# A proline-rich-domain-binding single domain antibody selectively inhibits RNA-induced liquid-liquid phase separation of tau

**DOI:** 10.64898/2026.01.09.698559

**Authors:** Simon Thiou, Leslie Martin, Evangelia Manousaki, Marine Nguyen, Justine Mortelecque, Leila Heidsieck, François-Xavier Cantrelle, David Blum, Valérie Buée-Scherrer, Luc Buée, Isabelle Landrieu, Elian Dupré, Clément Danis

## Abstract

Liquid-liquid phase separation (LLPS) mediates the formation of biomolecular condensates, which organize cellular processes such as synaptic plasticity and stress response. The neuronal microtubule-associated protein tau undergoes LLPS under specific conditions, regulating synaptic vesicle clustering and microtubule dynamics.

*In vitro*, tau LLPS is induced by cofactors such as polyethylene glycol (PEG) or RNA, mainly via weak multivalent electrostatic interactions. However, the molecular mechanisms governing the formation of tau LLPS, including domain specific contribution, remain unclear.

In this study, we used eight single-domain antibodies (VHHs), targeting six distinct short sequences of tau, to explore the mechanisms of tau LLPS *in vitro*. By combining several biophysical methods, we evaluated the effect of each anti-tau VHH on tau LLPS with two main LLPS inducers, PEG (molecular crowding) and RNA (complex coacervation).

With PEG as an inducer, all VHHs targeting tau enhanced tau LLPS formation, regardless of their affinity for tau. With RNA as an inducer, the effect of the VHHs was mixed: VHHs targeting the C-terminal domain promoted condensation, while VHH B1-1, which binds the proline-rich domain (PRD; including residues (_221_REPKKVAVVRTP_232_), abolished droplet formation. NMR and surface plasmon resonance confirmed 1 to 1 binding of VHH B1-1 to the PRD, and competition assays with a PRD peptide restored LLPS, demonstrating mechanistic specificity.

This result underscores the importance of this region in tau LLPS formation. Our findings provide domain-resolved insights into the regulation of tau LLPS and demonstrate the potential of VHHs as tools to selectively modulate biomolecular condensates in physiological and pathological contexts.

## Introduction

Liquid-liquid phase separation (LLPS) is a fundamental biophysical process in which a homogeneous solution, composed of proteins or nucleic acids, separates into two distinct liquid phases. LLPS forms through multivalent and weak interactions that sequester proteins (with or without nucleic acids) into liquid, membrane-less droplets. Intrinsically disordered proteins are the primary protein component of these phase separations due to their ability to engage in these types of molecular interactions. LLPS enables cells to rapidly adapt to change of environmental cues by concentrating or excluding specific biomolecules, thereby enabling the formation of subcellular structures called biomolecular condensates. LLPS plays important roles in organizing cellular components, regulating signaling pathways, and maintaining cellular homeostasis. In neurons, LLPS is a key mechanism in the regulation of synaptic plasticity, RNA metabolism, and stress responses.^1,2^

Tau is an intrinsically disordered, microtubule-associated neuronal protein that modulates axonal transport and plays multiple other roles in cells.^3^ Tau can undergo LLPS, and tau LLPS has been observed in the intracellular compartment of neuronal cells.^4,5^ Tau LLPS, also called tau condensates or tau droplets, seems to play a significant role in regulating microtubule formation.^6,7^ Tau droplets are also localized in the synaptic compartments, where they may organize presynaptic vesicles and contribute to neuronal communication.^8^ In addition, various biomolecules that interact with tau modulate tau LLPS. This includes interaction with chaperone-like proteins, which decrease tau phase separation, and interaction with RNA-binding proteins, such as Tia1 or TDP43, which facilitate it.^9–12^

Tau LLPS is mainly studied *in vitro* using two types of inducers (or combinations thereof) with different modes of action. Tau LLPS can be initiated by adding macromolecular crowding and dehydrating agents, such as polyethylene glycol (PEG).^4^ PEG excluding volume and solvation effects favor attractive forces between tau molecules, triggering the de-mixing of tau into liquid condensates. Tau LLPS can also be induced by a coacervation mechanism, whereby oppositely charged between tau and RNA spontaneously organize into higher order liquid condensates.^13,14^ These two processes are mainly driven by the electrostatic intra- and inter-molecular interactions of the tau protein. However, the molecular determinants of tau LLPS remain unclear. Thus far, they have been investigated *in vitro* using mutations, truncated constructs and domain deletions.^12,15,16^ It remains unclear which tau domain(s) contribute(s) to tau droplet formation under different LLPS-inducing conditions. Additionally, the interplay between tau and its binding partners, or tau post-translational modifications (PTMs), in regulating LLPS is not fully characterized.

To identify domain-specific regulators of tau condensation, we here tested a panel of eight single-domain antibodies, also called VHHs,^17^ which target six distinct short regions of tau (Figure 1).^18–20^ Their small size and high specificity allow for the precise modulation of tau’s conformational and interaction ensemble without disrupting its global structure. Using this panel, we systematically investigated the effect of targeting different epitopes on tau LLPS under two induction conditions: PEG and RNA. We also used the ability to chemically label the anti-tau VHHs with fluorophores to track tau LLPS formation. Using biophysical assays such as optical density (OD), dynamic light scattering (DLS), and fluorescence microscopy, we showed that, under molecular crowding conditions, all anti-tau VHHs enhanced tau’s ability to form LLPS, with the VHHs localizing inside the tau droplets. Under RNA coacervation conditions, anti-tau VHHs targeting the C-terminal domain of tau also appeared to increase tau droplet formation. However, a VHH targeting a positively charged epitope in the PRD domain of tau fully inhibited their formation. These findings suggest that interactions mediated through the PRD, rather than the microtubule-binding repeat domain, are crucial for the multivalent contacts that drive RNA-induced tau phase separation.

**Figure 1:**
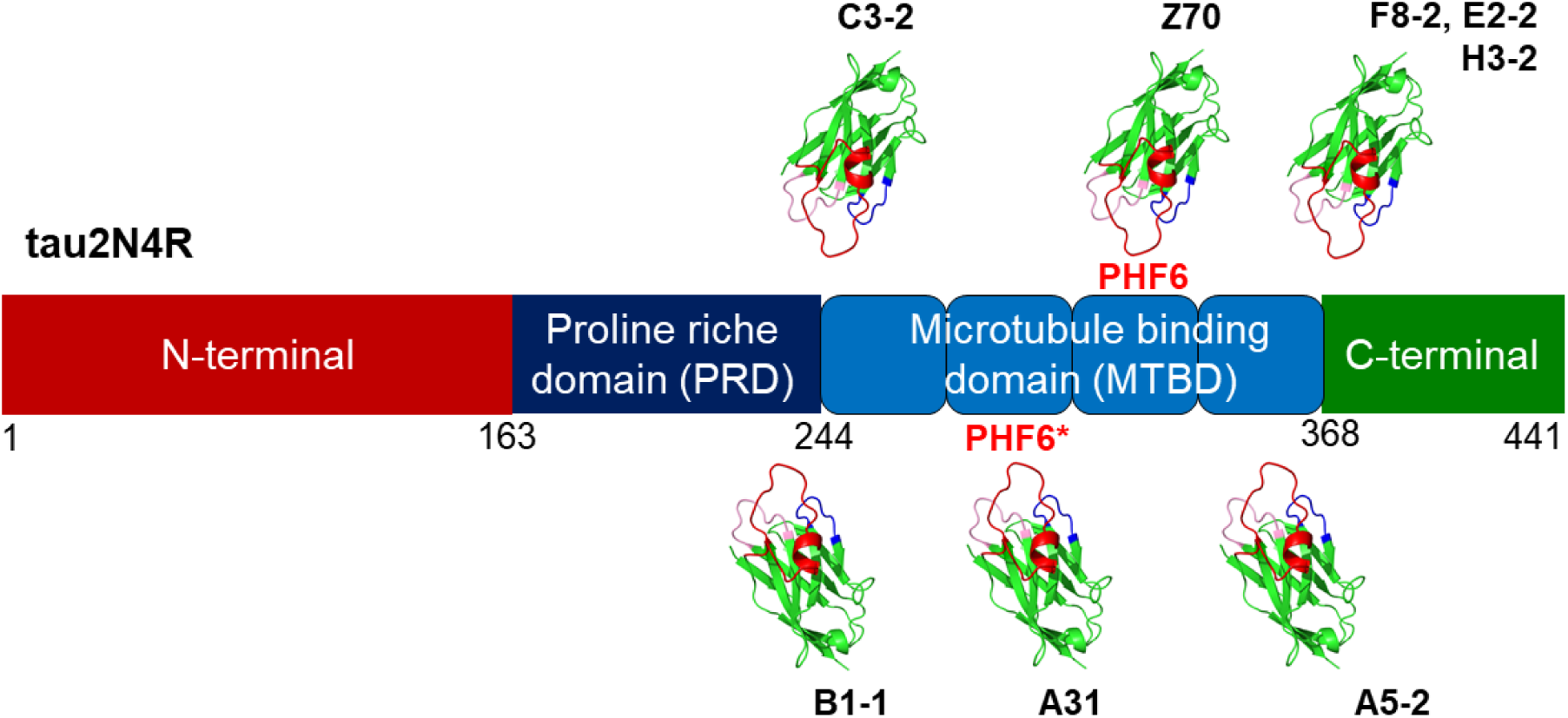
Overview of the tau epitopes recognized by the anti-tau VHHs used in this study and schematic representation of the different tau LLPS characterization assays. Schematic representation of the tau epitopes recognized by each anti-tau VHH. VHH B1-1 binds to the tau proline-rich domain (PRD). VHH C3-2 binds to an amino acid sequence located between the PRD and the R1 repeat of the tau microtubule-binding domain (MTBD). VHH A31 binds to the PHF6* region within the R2 repeat. VHH Z70 binds to the PHF6 region within the R3 repeat. VHH A5-2 binds to the R4 repeat. VHH F8-2, E2-2 and H3-2 bind to the C-terminal domain of tau. The epitopes were determined by 2D NMR interaction mapping experiments, as previously described.^19,29,35^

Together, these results demonstrate that the tau interaction network changes under different LLPS-inducing conditions, though electrostatic forces remain the dominant factor. This work provides domain-resolved insights into tau LLPS and introduces VHHs as probes that can selectively modulate biomolecular condensates. This has implications for understanding and potentially targeting tau mis-regulation in neurodegenerative diseases.

## Results

### Anti-tau VHHs increase tau condensation in molecular crowding conditions

We first investigated conditions that induce tau LLPS using polyethylene glycol (PEG) by monitoring the optical density at 400 nm (OD_400_). We incubated 10 μM of tau2N4R in a Tris buffer (50 mM Tris, 10 mM NaCl, pH 7.4) with low ionic strength in the presence of increasing concentration of PEG (10kDa). Under these conditions, the signal began to increase at 4% PEG and reached a plateau at 12% (Figure 2a). The concentration of PEG needed to reach 50 % of the maximal turbidity signal corresponded to 6.5% (Figure 2a). Phase diagram analysis confirmed that the increase in turbidity was dependent on the increase in PEG and tau concentrations (Figure 2b, Supplementary Figure 2a). To study the effect of anti-tau VHHs on tau LLPS formation, we selected a 7% PEG condition, which allows to observe any potential inhibitory or enhancing effects of the VHHs. We then incubated tau in the absence or presence of each anti-tau VHH at a 1:1 molar ratio (at 10 μM). We induced tau droplet formation by adding 7% of PEG to the mixture (tau LLPS, normalized to an OD_400_ of 1; Figure 2c). As control, we used a VHH targeting the V5 tag (anti-V5)^21^ that should not interfere with tau droplet formation because its epitope is absent and it does not bind to tau2N4R (Supplementary Figure 1). Additionally, tau LLPS was assayed in the presence of 300 mM of NaCl (NaCl), an inhibitor of tau droplet formation.^15^ As expected, we observed no change in the absorbance values when tau droplets were induced in the presence of anti-V5 (OD_400_ = 1.07 ± 0.06), whereas the presence of 300 mM NaCl fully inhibited tau droplet formation (OD_400_ = 0.04 ± 0.04) (Figure 2c). Interestingly, we observed an increase in OD_400_ values compared to tau LLPS with all the anti-tau VHHs used in this study, and especially in the presence of VHH Z70, F8-2, and E2-2, with increases of more than 1.5-fold (OD_400_ = 1.84 ± 0.38, 1.73 ± 0.25, and 1.76 ± 0.25, respectively) (Figure 2b). Next, we selected VHH F8-2 and performed an experiment in which we fixed its concentration at 10 µM while varying the concentrations of PEG and tau (Figure 2d, Supplementary Figure 2b). In the presence of VHH F8-2, the OD_400_ signal increased under all conditions beginning at 4μM of tau and 6-8% of PEG (Figure 2d, Supplementary Figure 2b). Tau LLPS formation was further visualized using labeled tau-TAMRA confocal microscopy analysis (Figure 2e-f; Supplementary Figure 3), in all the conditions except in the presence of 300 mM NaCl (Supplementary Figure 3).

**Figure 2:**
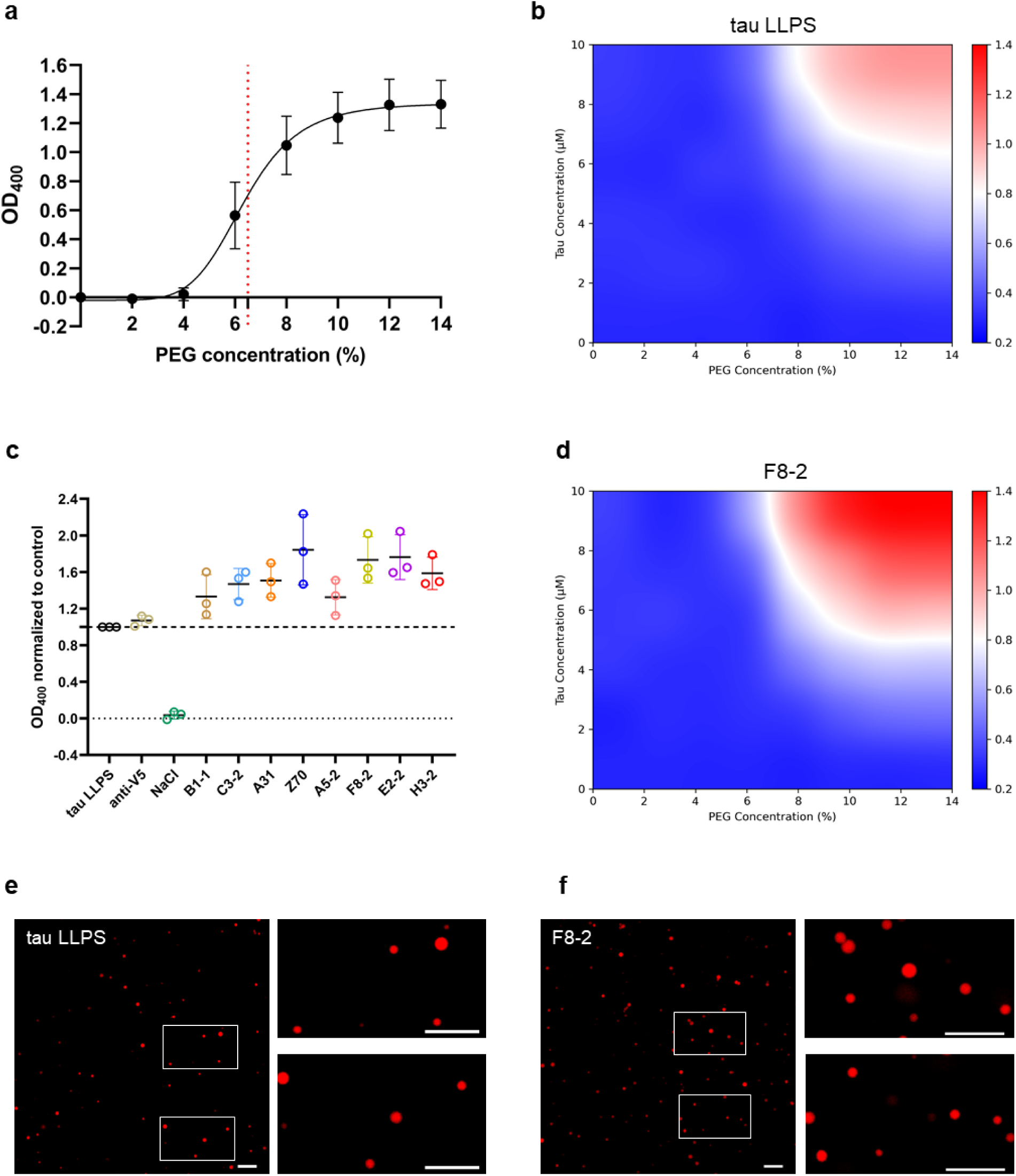
Tau co-incubated with anti-tau VHHs increases tau LLPS formation in the presence of PEG **a.** The concentration response curve shows the effect of different PEG concentrations (expressed as percentages) on 10 μM tau2N4R LLPS. The horizontal dashed line corresponds to the concentration of PEG needed to reach 50 % of the maximal signal of OD_400_ **b.** Phase diagram of tau2N4R. **c.** The OD_400_ value corresponding to tau LLPS formation in the presence of 10 μM tau2N4R and 7% PEG (tau LLPS) was normalized to 1. Under the same conditions, tau2N4R was pre-incubated with either 10 μM of an anti-V5 VHH (anti-V5), 300 mM NaCl (NaCl), or 10 μM of each anti-tau VHH. Three independent experiments were performed at least in duplicate for each condition. The scatterplot representation includes the mean and standard deviation (SD). **d.** Phase diagram of tau2N4R in the presence of VHH F8-2 at a fixed concentration of 10 μM. The color coding represents the average optical density value at 400 nm. The color coding represents the average optical density value at 400 nm. **e.** Representative fluorescence image and associated zoom images of tau LLPS formed in the presence of 10 μM of tau-TAMRA and 7% of PEG. Scale bar: 10 µm. **f.** Representative fluorescence image, and associated enlarged images, of tau LLPS formed in the presence of 10 μM tau-TAMRA, which was co-incubated with 10 μM anti-tau VHH F8-2 and 7% of PEG. Scale bar: 10 µm.

We analyzed the particle size distribution of tau under various conditions using dynamic light scattering (DLS) (Figure 3a). In the absence of PEG, the measurements confirmed the presence of particles primarily corresponding to tau monomers with a hydrodynamic radius (R_h_) of 6.2 ± 0.3 nm, consistent with previous observations. Larger size particles were also observed and could correspond to tau multimers.^22,23^ With the addition of 7% PEG (tau LLPS), we observed the formation of tau droplets (R_h_: ∼ 1500-2000 nm) (Figure 3a). Adding 300 mM of NaCl inhibited the formation of tau droplets as no particles of size with a R_h_ higher than 500 nm were detected. The presence of anti-V5 or anti-tau VHHs led to the formation of droplets with a similar size distribution to that of the tau LLPS condition (Figure 3a), suggesting that the overall size distribution of tau LLPS are not affected by the presence of the anti-tau VHHs. Using fluorescence recovery after photobleaching (FRAP), the tau droplets exhibited fluorescence recovery consistent with the liquid-like internal mobility of the tau-TAMRA droplets (Figure 3b-c). Even in the presence of VHH F8-2, which strongly affected the OD_400_ signal, no difference in fluorescence recovery was observed compared to the control condition (Figure 3b-c), suggesting that VHH F8-2 does not influence the liquid state of the tau LLPS.

**Figure 3:**
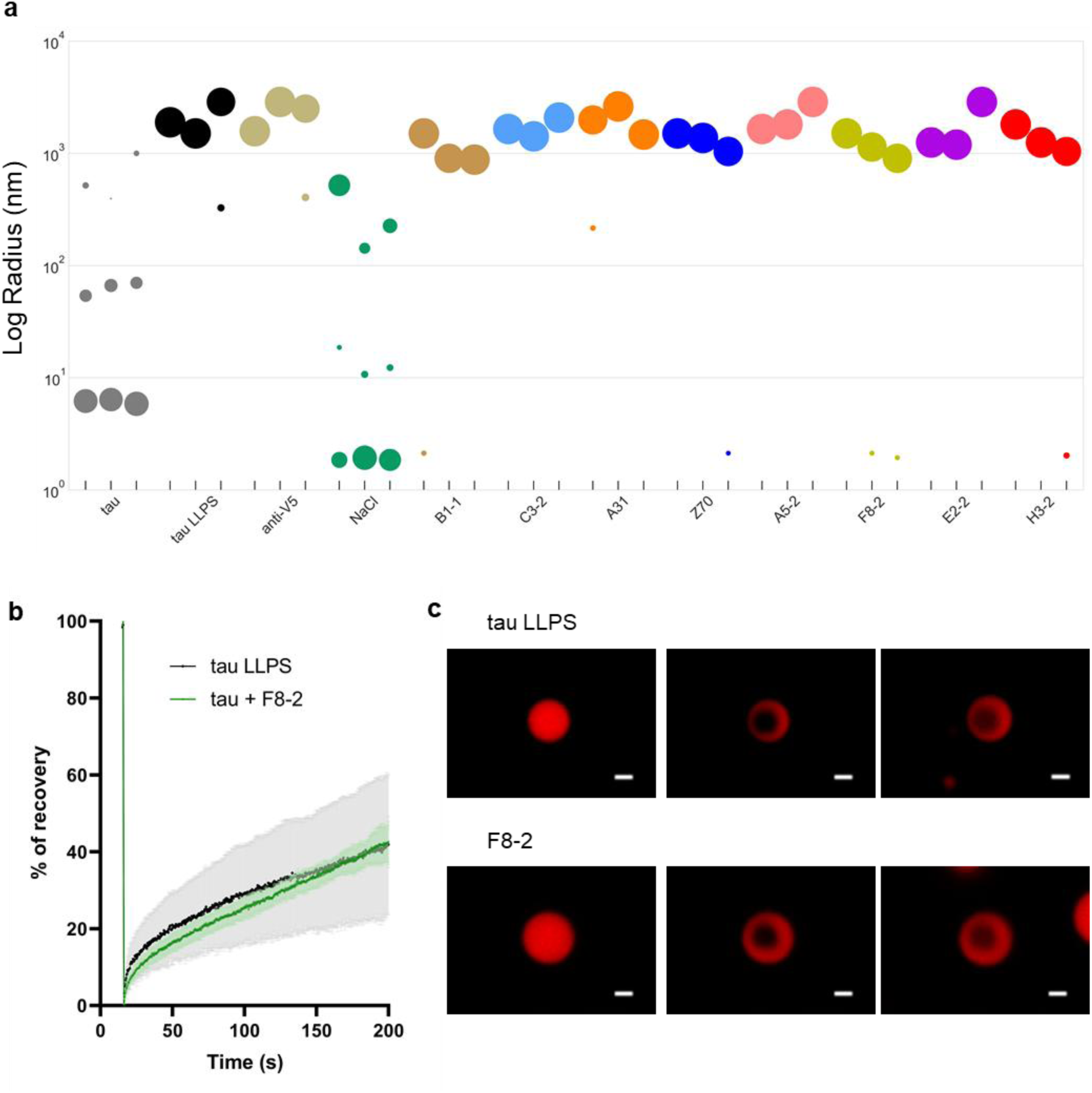
PEG-Tau LLPS pre-formed with anti-tau VHHs visualized by fluorescent microscopy did not modify the size distribution and the liquid-state of tau droplets. **a.** Representative DLS detection of 10 μM fixed concentration of tau in solution without PEG (tau), with 7% of PEG (tau LLPS), pre-incubated with 10 μM of a VHH targeting the V5 tag (anti-V5), 300 mM NaCl (NaCl) or 10 μM of each anti-tau VHH. The data correspond to three independent experiments, each with ten data points. The size of the spheres represents the intensity of the scattered light from the particles detected in solution. **b**. Fluorescence recovery after photobleaching (FRAP) measurements of tau-TAMRA (10 μM) droplets formed in the absence (black curve) or presence (light green curve) of 10 μM VHH F8-2 after LLPS induction with 7% of PEG. **c.** Representative images of the FRAP recovery of tau-TAMRA droplets (pre-bleach T15s, post-bleach T16s and post-bleach T200s) in the absence (top panels) or presence (bottom panels) of VHH F8-2 illustrating the fluorescence recovery after photo-bleaching. Scale bar is 1 µm.

In summary, these experiments demonstrated that the formation of tau LLPS in molecular crowding conditions is enhanced by the presence of anti-tau VHHs that bind to distinct epitopes as demonstrated by the increasing turbidity signal, although the size distribution and the liquid-state of tau droplets were not affected.

### Anti-tau VHHs that increase tau-PEG droplets formation are located inside the tau droplets

To better characterize the effect of the anti-tau VHHs on tau droplet formation, we quantified the labeled surface area, which depends on both the number and the size of tau droplet, using fluorescence confocal microscopy analysis (Supplementary Figure 4). Tau droplet formation was more extensive in the presence of most anti-tau VHHs compared to the tau LLPS and anti-V5 conditions (area = 18.8 and 24.5 μm^2^, respectively), with the strongest effects observed with VHHs Z70, A31, E2-2, and F8-2 (area = 40.7, 36.3, 31 and 30.3 μm2, respectively) (Figure 4a). As expected, there was a strong correlation between the *in vitro* OD_400_ assay and the labeled surface area (Pearson correlation coefficient of 0.93; Figure 4b). The VHHs with the highest values in the turbidity assay clearly produced a higher number and/or size of tau droplets, demonstrating a clear link between the turbidity signal and the presence of tau droplet. Since the VHHs were selected for their ability to bind specifically to tau, we hypothesized that the direct binding of the VHHs to tau caused the increased propensity of tau to undergo LLPS. Thus, we performed a fluorescence microscopy experiment using labeled tau and VHHs with different fluorescent dyes. We prepared the labeled tau LLPS *in vitro* in the presence of each of the two VHHs showing the strongest effects on tau droplet formation: Z70 and F8-2. Thanks to this orthogonal labeling strategy, we observed that both VHHs are localized inside tau droplets (Figure 4c, Supplementary Figure 5), which strongly suggests that their direct interaction increases tau droplet formation.

**Figure 4:**
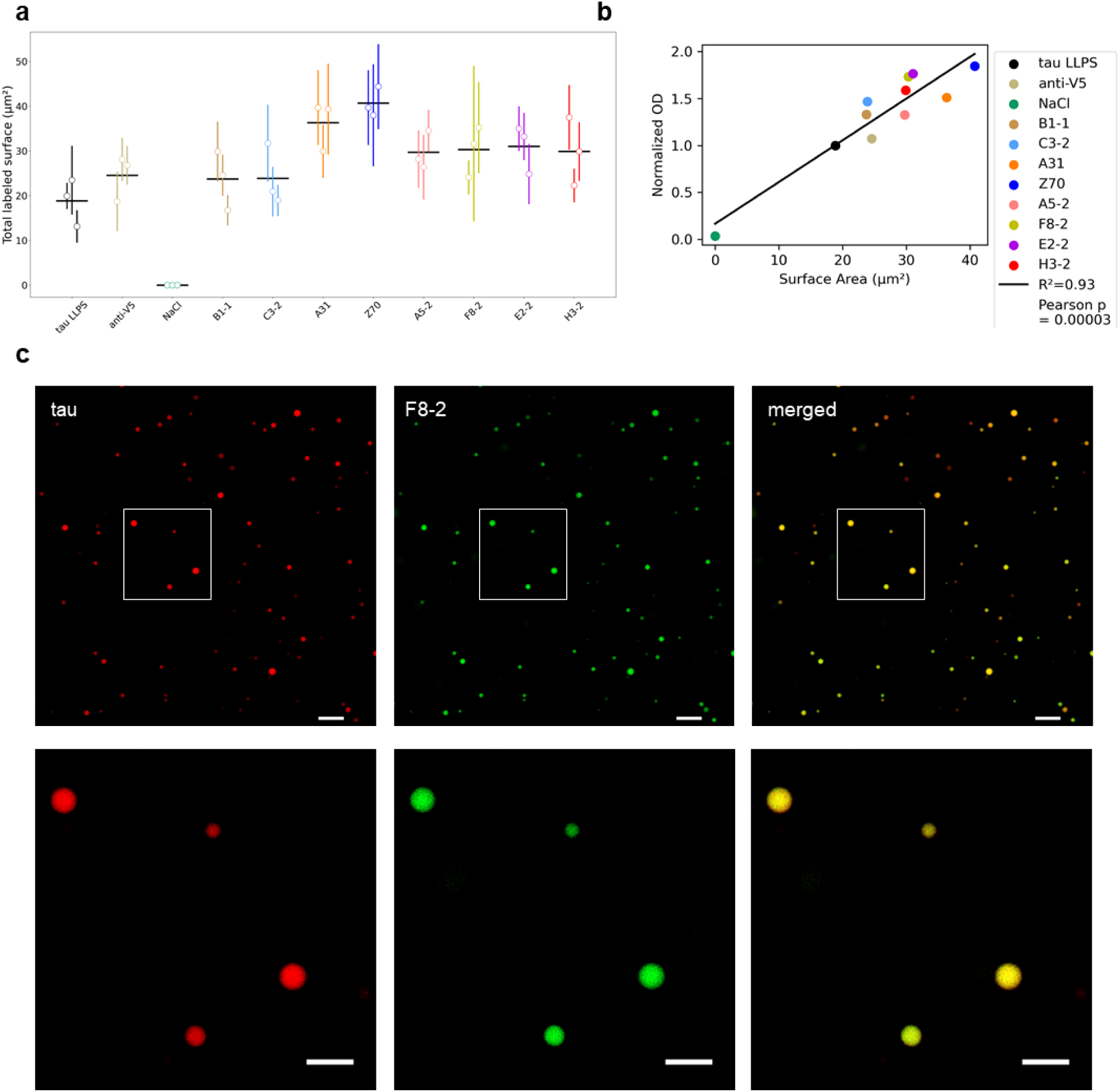
Anti-tau VHHs increase the formation of PEG-tau droplets and are located inside the droplets. **a.** Quantification of the total labeled surface area is based on the TAMRA fluorescence signal under conditions resulting from tau LLPS formed in the presence of 10 μM of tau-TAMRA and 7% of PEG (tau LLPS), which was pre-incubated with either 10 μM of an anti-V5 VHH (anti-V5), 300 mM NaCl (NaCl), or 10 μM of each anti-tau VHH. Data are presented as the mean of three independent replicates per condition, along with the associated standard error. **b.** The Pearson plot shows the total labeled area of the tau droplets versus the normalized optical density values obtained in Figure 2b. **c.** Representative fluorescence images, and associated enlarged images of tau droplets, formed in the presence of 10 μM of tau-TAMRA, which was co-incubated with 10 μM of VHH F8-2-Atto488 and 7% of PEG. Scale bars are 10 µm and 5 µm, respectively.

### Anti-tau VHHs differentially modulate tau LLPS formation under complex coacervation conditions

Nucleic acids, such as RNA, are known to be involved in forming condensates that regulate various intracellular functions. Since RNA may be a more physiologically relevant inducer of tau LLPS than PEG, we investigated the modulation effect of the anti-tau VHHs on tau LLPS in the presence of RNA.^14^ We thus incubated 20 μM of tau2N4R with increasing concentrations of RNA polyA in a HEPES buffer at low ionic strength and monitored the turbidity changes (OD_400_). Under these conditions, the OD_400_ signal indicative of tau droplet formation increased between 3 and 10 μg/mL of RNA and reached a plateau at 100 μg/mL (Figure 5a). Concentration of RNA needed to reach 50 % of the maximal signal of turbidity corresponded to 55 μg/mL (Figure 5a). We selected the condition of 20 μM of tau and 50 μg/mL of RNA to evaluate the VHHs’ effect. We then incubated tau in the absence or presence of each anti-tau VHH using a 1:1molar ratio, at 20 μM. We induced the formation of tau droplets by adding 50 μg/mL of RNA to the mixture (tau LLPS, normalized to an OD_400_ of 1; Figure 5b). Similar to molecular crowding conditions, the OD_400_ value remained unaffected when tau droplets were induced in the presence of the anti-V5 VHH (OD_400_ = 0.93 ± 0.11). However, the presence of 300 mM NaCl fully inhibited the formation of tau droplets (OD_400_ = 0.03 ± 0.07). We observed an increase in the OD_400_ value compared to the tau LLPS control, primarily in the presence of VHH F8-2, which targets the tau C-terminal epitope and increased the signal by 30 % (1.30 ± 0.13) (Figure 5b). Conversely, we found that other VHHs tended to decrease the OD_400_ values. Specifically, VHH B1-1 had an inhibitory activity (OD_400_ = 0.13 ± 0.18, Figure 5b). Additionally, we established a phase diagram illustrating the effect of different concentrations of RNA (ranging from 0 to 125 μg/mL) and tau (ranging from 0 to 20 μM) in the absence or in the presence of a fixed concentration of VHH B1-1 (20 μM) (Figure 5c-d, Supplementary Figure 6a-b). Phase diagram analysis confirmed that the increase in turbidity signal was depending on the increase in the tau and RNA concentration. However, we also observed that strong RNA concentration in the presence of weaker tau concentration decrease turbidity signal (Figure 5c, Supplementary Figure 6a). Analysis of the phase diagram confirmed a significant decrease in turbidity value in the presence of VHH B1-1 at all tested concentrations of RNA and tau (Figure 5d, Supplementary Figure 6b). We analyzed the size distribution of tau-VHH droplets in solution Using DLS (Figure 5e). These measurements confirmed the major presence of particles corresponding to tau monomers with an R_h_ of 5.7 ± 0.1 nm, which is consistent with previous observations. The presence of larger particles, as observed in Tris buffer, is consistent with the formation of soluble multimers.^22,24^ After adding RNA (tau LLPS), we observed the formation of tau droplets (R_h_ ∼ 450 nm) (Figure 5e). The presence of anti-V5 or anti-tau VHHs (C3-2, A31, Z70, A5-2, F8-2, E2-2 and H3-2) did not strongly affect the particle size distribution, compared to tau alone (R_h_ ∼ 200 - 700 nm, Figure 5e), suggesting that the overall size distribution of tau LLPS was unaffected by the presence of these anti-tau VHHs. Adding 300 mM NaCl inhibited tau droplet formation as almost no particles with an R_h_ greater than 200 nm were detected, and particles similar in size to monomeric tau were observed (R_h_ of 7.2 ± 0.4 nm). Interestingly, the size distribution of tau droplets was also disrupted in the presence of VHH B1-1, with smaller particles detected (R_h_ of 24.4 ± 1.9 nm, Figure 5e).

**Figure 5:**
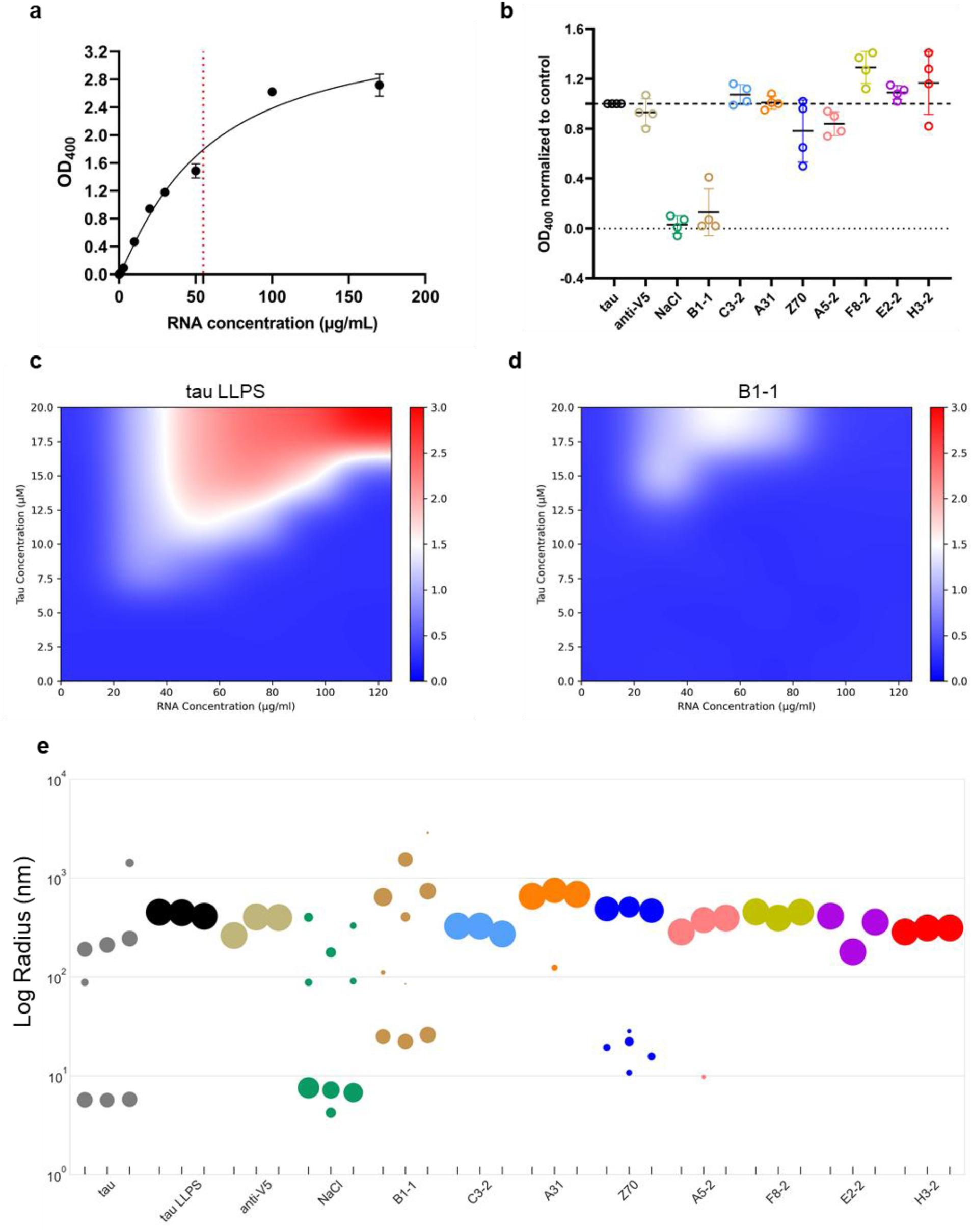
Co-incubation with anti-tau VHHs increases or decreases the turbidity associated with tau LLPS formation in the presence of RNA. **a.** The concentration response curve of RNA polyA (in μg/mL) is shown in the presence of a fixed concentration of 20 μM tau2N4R. Fifty percent of the maximal OD_400_ value is reached at a concentration of 50 μg/mL RNA polyA. The horizontal dashed line corresponds to the concentration of RNA needed to reach 50 % of the maximal signal of OD_400_ **b.** The normalized OD_400_ values correspond to tau LLPS formation in the presence of 20 μM of tau2N4R and 50 μg/mL of RNA (tau LLPS), pre-incubated with either 20 μM of an anti-V5 VHH (anti-V5), 300 mM NaCl (NaCl), or 20 μM of each anti-tau VHH. Four independent experiments were performed at least in duplicate for each condition. Data were normalized to the tau LLPS condition. The scatterplot includes the mean and standard deviation (SD). **c.** Phase diagram of tau2N4R. **d.** Phase diagram of tau2N4R in the presence (right) of VHH B1-1 (fixed concentration of 20 μM). Color coding represents the average optical density value at 400 nm. **e.** Representative DLS detection of a 20 μM fixed concentration of tau in solution without RNA polyA (tau), with 50 μg/mL of RNA polyA (tau LLPS), pre-incubated with 20 μM of a VHH targeting the V5 tag (anti-V5), 300 mM NaCl (NaCl), or 20 μM of each anti-tau VHH. Three independent experiments were performed, each with ten data points. The size of the spheres represents the intensity of the scattered light of the respective particles detected in solution.

We further confirmed tau LLPS formation using labeled tau-TAMRA confocal microscopy analysis (Figure 6a-b; Supplementary Figure 7). We visualized tau droplets in all of our conditions except in the presence of 300 mM NaCl and VHH B1-1, in which few or no tau droplets were detected (Supplementary Figure 7). We quantified the labeled surface area, in our different conditions, using fluorescence confocal microscopy analysis. Although there was more variability among the experiments than with the PEG ones, we observed an increase in the number and/or the size of tau droplets with VHH F8-2 compared to the tau LLPS condition (area = 370 and 185 μm^2^, respectively, Figure 6c). Conversely, we observed a significant decrease in tau droplet formation in the presence of VHH B1-1 (area = 8 μm^2^, Figure 6c). The correlation between the *in vitro* OD_400_ values and the labeled surface area (Pearson correlation coefficient of 0.78; Figure 6d) confirmed that the turbidity signal monitored tau droplet formation. Altogether, these experiments demonstrate that the formation of tau LLPS in coacervation conditions is enhanced by the presence of anti-tau VHH F8-2 and is inhibited by the presence of anti-tau VHH B1-1.

**Figure 6:**
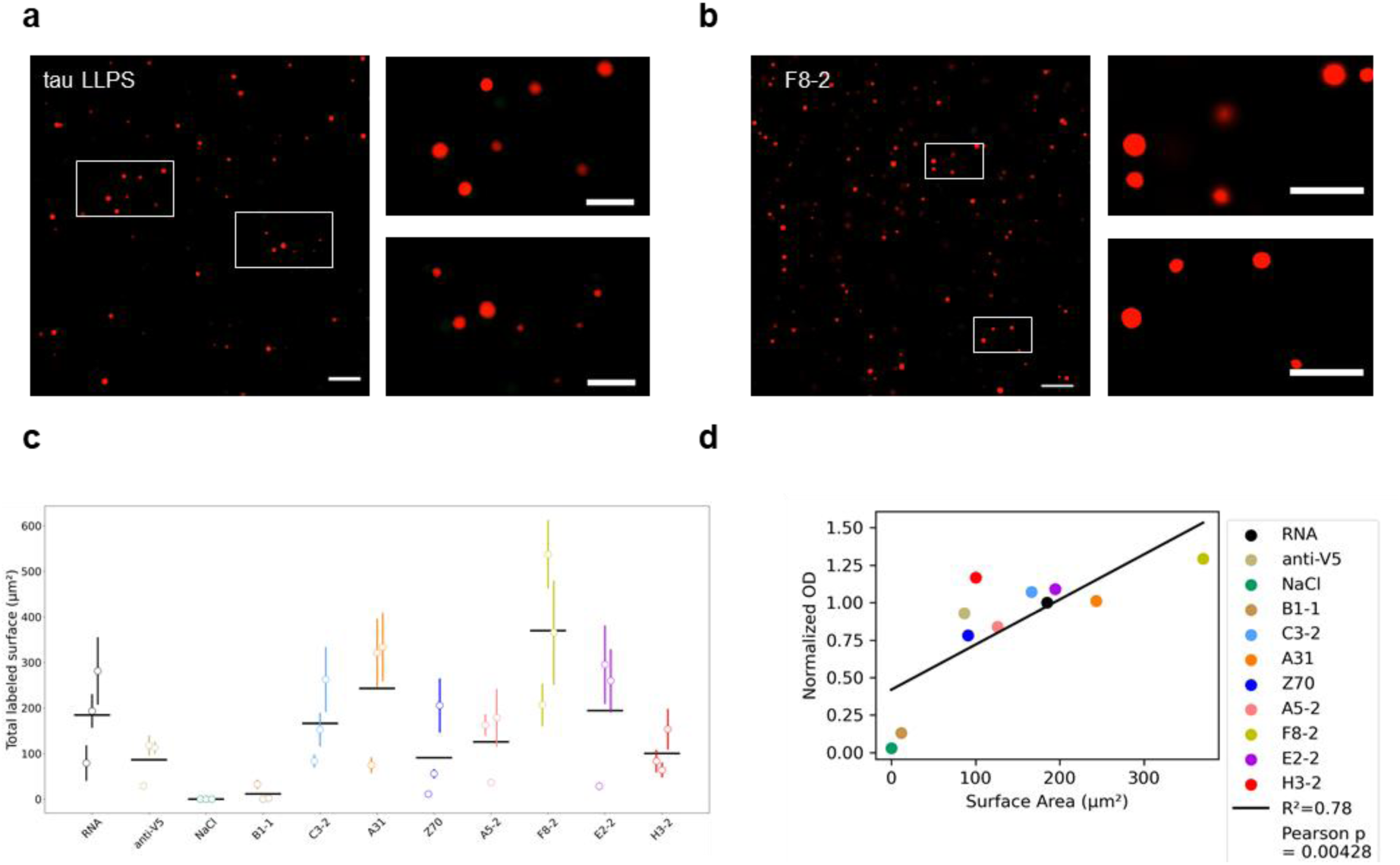
RNA-Tau LLPS pre-formed with anti-tau VHHs is associated to an increase or decrease of tau droplets **a.** Representative fluorescence image, and associated enlarged images, of tau droplets formed in the presence of 20 μM of tau-TAMRA and 50 μg/mL of RNA polyA. Scale bar: 20 µm and 10 µm respectively. **b.** Representative fluorescence images, and associated enlarged images, of tau droplets formed in the presence of 20 μM of tau-TAMRA co-incubated with 20 μM of anti-tau VHH F8-2 and 50 μg/mL of RNA polyA. Scale bar: 20 µm and 10 µm respectively. **c.** Quantification of the total labeled area of the tau droplets, based on the TAMRA fluorescence signal, in conditions resulting in tau LLPS. This was formed in the presence of 20 μM of tau-TAMRA and 50 μg/mL of RNA polyA(tau LLPS). It was pre-incubated with either 20 μM of an anti-V5 VHH (anti-V5), 300 mM NaCl (NaCl), or 20 μM of each anti-tau VHH. Data are presented as the mean of three independent replicates per condition, along with the associated standard error. **d.** The Pearson plot shows the total labeled area of the tau droplets versus the normalized optical density values obtained in Figure 5b.

### The anti-tau VHH F8-2 enhances the formation of tau RNA droplets and is located inside the droplets

To better understand the mode of action of VHH F8-2, which had the strongest effect on tau droplet formation in the presence of RNA, we performed fluorescence microscopy experiments. Tau and VHH F8-2 were labeled with the fluorophores TAMRA and Atto488, respectively. Our results showed that VHH F8-2 was localized inside the tau droplets (Figure 7a), suggesting a direct interaction that increases tau droplet formation. Using FRAP, we confirmed the liquid-like state of tau-TAMRA LLPS (Figure 7b-c). The presence of VHH F8-2, did not strongly affect the fluorescence recovery compared to the control condition (Figure 7b-c). In summary, these experiments suggested that the formation of tau LLPS in coacervation conditions is enhanced by the direct binding of VHH F8-2 to tau but did not strongly affect the liquid state of tau droplets.

**Figure 7:**
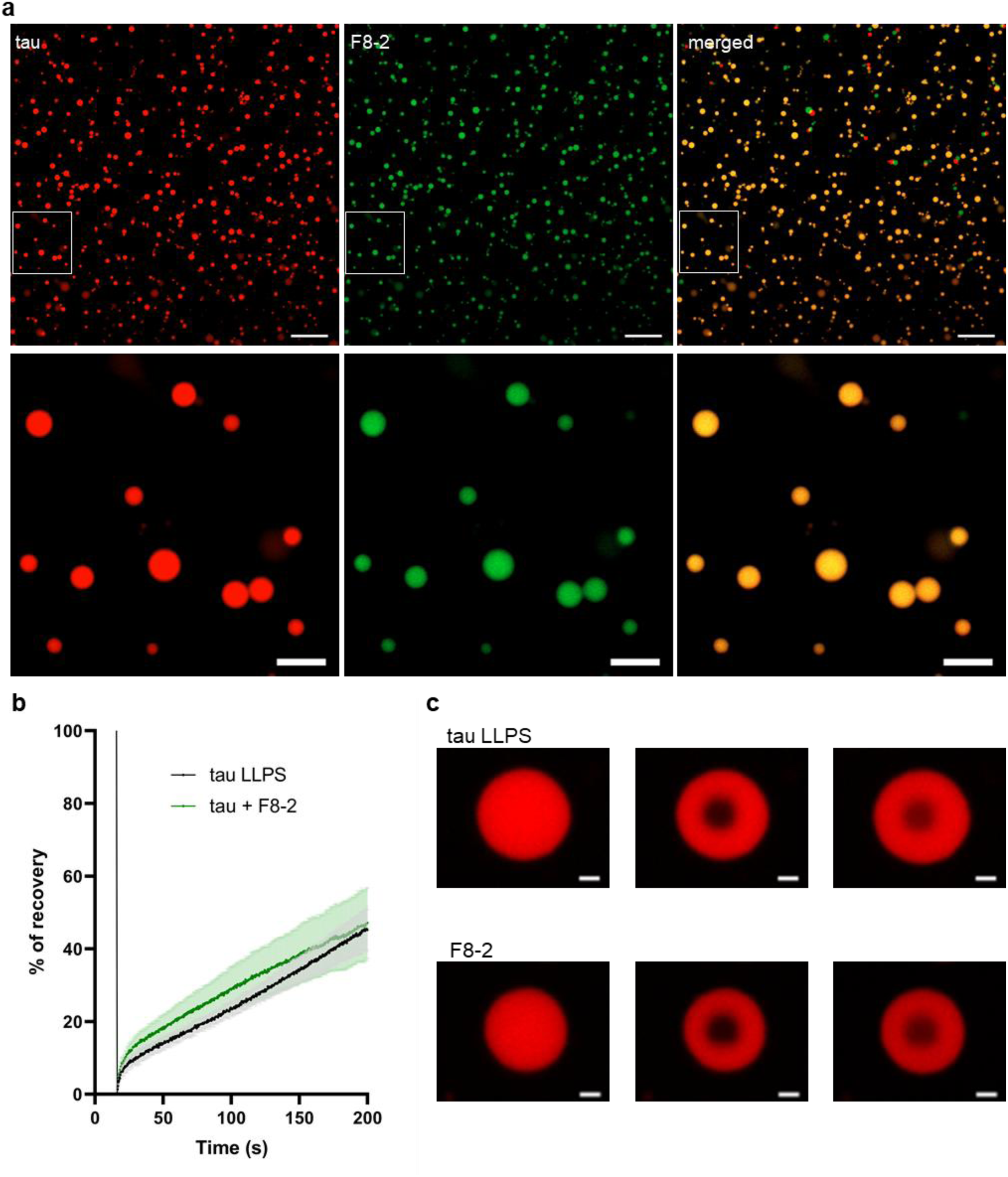
Anti-tau VHHs F8-2 is located inside the tau droplets and modify liquid transition state of tau droplets. **a.** Representative fluorescence image and associated zoom images of tau LLPS formed in the presence of 20 μM tau-TAMRA co-incubated with 20 μM VHH F8-2-Atto488 and 50 μg/mL. Scale bar: 20 µm and 5 µm, respectively. **b**. Fluorescence recovery after photobleaching (FRAP) measurements of tau-TAMRA (20 μM) droplets formed in the absence (black curve) or in presence (light green curve) of 20 μM VHH F8-2 after LLPS induction with 50 μg/mL. **c.** Representative FRAP images of tau-TAMRA droplets (pre-bleach T15s, post-bleach T16s and post-bleach T200s) in the absence (top panels) or presence (bottom panels) of VHH F8-2., illustrating the fluorescence recovery after photo-bleaching. Scale bar is 1 µm.

### The binding of anti-tau VHH B1-1 to its proline-rich domain epitope inhibits tau LLPS in complex coacervation conditions

Since VHH B1-1 is the only VHH in our panel that strongly inhibits tau-RNA droplet formation, we next evaluated its mechanism of action. We previously reported that VHH B1-1 binds to a tau epitope located in the PRD domain.^19^ However, these studies were performed using the VHH tagged with a negatively charged c-myc peptide attached at the C-terminal position. In this study, we performed additional experiments to refine the exact epitope. Since the tau LLPS studies were performed using recombinant VHHs lacking any tag, we conducted further NMR investigation to study the tau-VHH B1-1 interaction. Resonance perturbation mapping in the ^1^H, ^15^N HSQC spectra of ^15^N-Tau, obtained by nuclear magnetic resonance (NMR) spectroscopy, allowed to define the VHHs B1-1 binding site along tau sequence. Comparing the spectra of tau alone in solution or in the presence of a VHH B1-1 identified the spectral perturbations that are used to define the binding region (Supplementary Figure 8). B1-1 affected many resonances in the tau spectrum corresponding to residues within the PRD domain, as well as the R2-R3 repeats of the MTBD (Supplementary Figure 8). Binding mapping was improved and confirmed using a tau fragment called tauF3, which encompasses tau residues from 208 to 324. This tau fragment is smaller (116 amino acid residues instead of 441) resulting in fewer resonances and thus less resonance overlaps in the corresponding TauF3 ^1^H, ^15^N spectrum. This makes the binding region easier to identify (Supplementary Figure 9). The affected resonances corresponded to amino acid residues located in a peptide extending from residue 220 to 243. However, the R2-R3 repeat region in the F3 fragment was unaffected by VHH B1-1 binding (Supplementary Figure 9), unlike what was observed for tau. To confirm the minimal epitope sequence recognized by VHH B1-1, an epitope mapping was performed using the yeast two-hybrid system. A library of tau fragments (N-terminal GAL4-activation domain fusion,GAL4-AD-tau_fragments) served as preys, and VHH B1-1 (LexA C-terminal fusion, VHH-LexA fusion) served as bait. Twenty-four positive clones, indicative of VHH B1-1 binding to one tau fragment, were selected from a small-scale cell-to-cell mating screen. The minimal common amino acid sequence in all the corresponding tau fragment prey sequences was identified as peptide _221_REPKKVAVVRTPPKSPSSAKSRLQT_245_ of tau as the epitope of VHH B1-1 (Supplementary Figure 10). This result is consistent with the NMR data.

Based on this interaction data, we examined the binding of VHH B1-1 to a tau PRD peptide, which corresponds to tau site _221_REPKKVAVVRTP_232_ and showed the strongest NMR spectral perturbations (i.e. signal loss of the corresponding resonances). We used biotinylated VHH B1-1 immobilized on the surface of a streptavidin-functionalized chip and the tau PRD peptide as the analyte in multi-cycle kinetics (MCK), which consists in ten cycles. Each cycle corresponds to an injection of an increasing peptide concentration (Figure 8a). We determined the resulting dissociation constant K_D_ of the interaction using steady-state analysis by reporting the maximum binding response for each peptide concentration. The stoichiometric ratio (SR), which estimates the stoichiometry of an interaction, confirmed the 1 to 1 binding mode of the interaction (SR = 0.7). In this experimental setup, VHH B1-1 presented an affinity of 4.3 ± 1.3 μM, which confirmed its binding to this specific tau region (Figure 8b). Additionally, SPR was used to characterize the interaction between VHH B1-1 and a tau fragment corresponding to the MTBD of tau (tauMTBD) by immobilizing biotinylated tauMTBD on the surface of a streptavidin-functionalized chip (Supplementary Figure 10). Single-cycle kinetics (SCK) experiments confirmed the VHH B1-1 binding to the tau F3 fragment (K_D_ = 210 nM), consistent with the KD measured using tau2N4R (K_D_ = 289 nM).^20^ However, we did not detect any direct interaction between the tauMTBD fragment and the VHH B1-1. This finding is consistent with the tau epitope determined for VHH B1-1 (Supplementary Figure 11). Thus, the observed spectral perturbation of signals corresponding to the tauMTBD fragment in the full-length tau thus did not correspond to direct binding. Rather, we hypothesized that they could arise from long-range conformational effects.

**Figure 8:**
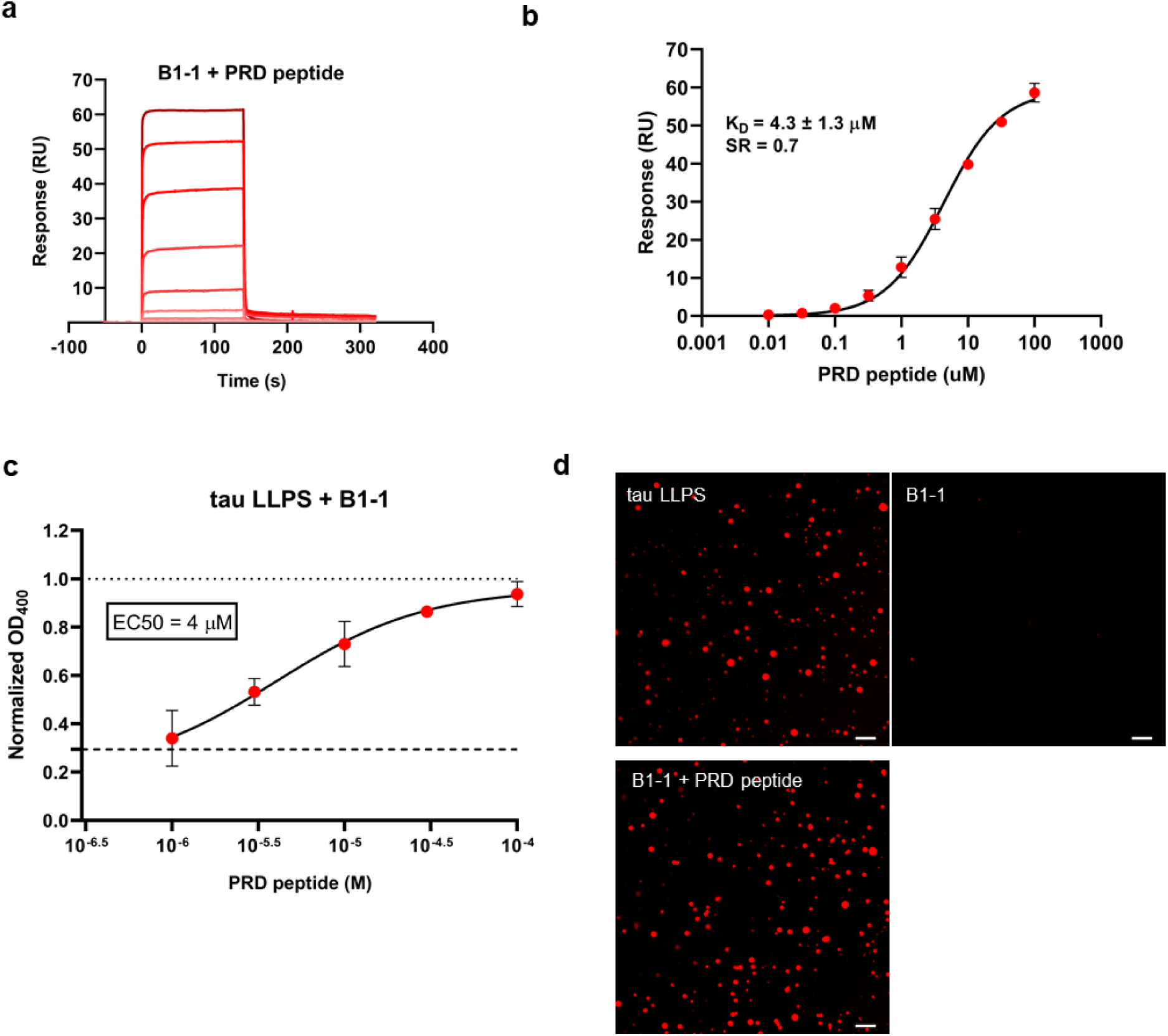
B1-1 binding to its proline rich domain epitope is responsible of tau LLPS inhibition in coacervation condition. **a.** Sensorgrams (reference-subtracted data) from multi-cycle kinetics (MCK) analyses performed on immobilized biotinylated VHH B1-1, with injections of increasing concentrations of the tau PRD peptide _221_REPKKVAVVRTP_232_ (0.5 log dilution, from 10 nM to 100 µM; n = 3 independent experiments). **b.** The maximum response (RU) observed for each peptide concentration was plotted and represented as a concentration-response curve (CRC). The black lines correspond to the fitted curves and the red dots correspond to the mean of the maximum response (in RU), for each peptide concentration. Error bars represent standard deviation. The K_D_ and stoichiometric ratio (SR) values extracted from these data are shown. K_D_s were obtained using the steady-state fitting model and are presented as mean values ± standard deviation. The SR was calculated from the estimated Rmax of three independent experiments. **c.** Concentration-response curve of the tau PRD peptide (0.5 log dilution, from 1 µM to 100 µM; n = 3 independent experiments) in the presence of a 20 μM fixed concentration of tau2N4R with 50 μg/mL of RNA, pre-incubated with 20 μM of VHH B1-1. The tau LLPS control condition corresponds to the dashed horizontal line (DO_400_ has been normalized to 1). Tau LLPS formed in the presence of VHH B1-1, but not the PRD peptide, correspond to the thick horizontal line. **d.** Representative fluorescence images tau droplets formed in the presence of 20 μM of tau-TAMRA and 50 μg/mL of RNA polyA (tau LLPS), in the presence of 20 μM VHH B1-1 (B1-1), and in the presence of 20 μM VHH B1-1 and 100 μM PRD peptide (B1-1 + PRD peptide). Scale bar: 10 µm.

To confirm that interaction with the VHH B1-1 to this tau PRD short motif was necessary to inhibit tau-RNA droplet formation, we conducted additional experiments, using the OD_400_ assay to monitor tau droplet formation. We incubated 20 μM of tau with a fixed concentration of 20 μM of VHH B1-1, as well as with increasing concentrations of the tau PRD peptide (ranging from 1 to 100 μM). We normalized the data to the condition in which tau droplets were formed with of 50 μg/mL of RNA (Figure 8c; dashed horizontal line). We validated the inhibition of tau droplet formation in the presence of VHH B1-1 (Figure 8c; dashed horizontal line OD_400_ = 0.29 ± 0.25). Interestingly, we observed an increase in the OD_400_ value as the concentration of the PRD peptide increased. The OD_400_ value reached a plateau at 100 μM of peptide (OD_400_ = 0.94 ± 0.05) (Figure 8c). We estimated the EC_50_ of the interaction and obtained a value close to the estimated K_D_ between VHH B1-1 and the PRD peptide (EC_50_ = 4 μM). Recovery of tau LLPS formation by the PRD peptide titration was confirmed by labeled tau-TAMRA confocal microscopy analysis (Figure 8d). We visualized the formation of tau droplets in the control condition (tau LLPS). VHH B1-1 inhibited their formation (B1-1), but the presence of 100 μM of PRD peptide (B1-1 + PRD peptide) restored it (Figure 8d).

We confirmed that VHH B1-1 binds to the tau PRD domain within the 220-243 region, and that tau LLPS inhibition depends on binding of VHH B1-1 to its epitope. Together, the NMR data, two-hybrid epitope mapping, and additional SPR experiments, which showed no direct binding of VHH B1-1 to the MTBR, suggest that the binding induces tau conformational rearrangement.

## Discussion

In this study, PEG and RNA were used as model inducers of tau LLPS, mimicking the crowded intracellular environment and the role of nucleic acids in cellular condensates, respectively.

Despite their short specific binding sites, we observed that single domain antibodies enhance tau LLPS in the presence of PEG. This effect cannot be attributed to a simple protein concentration effect, as the control VHH (anti-V5), which does not bind to tau, had no effect on tau LLPS. Notably, the affinity of the VHHs for tau does not appear to be a critical factor. For instance, VHHs E2-2 and F8-2, which recognize the same epitope with different binding affinities (distinct K_D_ values of 753 and 84 nM, respectively)^20^, both promote tau LLPS. VHH F8-2 was also found to co-localize with tau within the droplets, suggesting direct binding under LLPS conditions. The effect of the VHHs is epitope-dependent, with the strongest enhancement effect observed with VHHs binding the PHF6 or the C-terminal sequence of tau. On the other hand, other tau-binding proteins, such as holdase/chaperone proteins, inhibit tau LLPS in PEG conditions.^9^ However, these proteins bind to much larger regions of tau proteins through multiple weak interactions rather than presenting short, specific peptide epitopes, as VHHs do. To our knowledge, this type of monovalent, sequence-specific interaction with a short sequence, as observed with the VHHs here, has not yet been demonstrated to modulate tau LLPS, including the enhancement of PEG-induced LLPS.

Under RNA-induced conditions, the VHHs had a more pronounced and selective effect on tau LLPS. While some VHHs enhanced tau LLPS, others, most notably VHH B1-1, specifically inhibited tau droplet formation. VHH B1-1, which binds the epitope _221_REPKKVAVVRTPPKSPSSAKSRLQT_245_ in the PRD2 region of tau, selectively inhibited tau LLPS. This inhibitory effect is directly linked to this binding, as demonstrated by competition assays with a peptide mimicking the epitope. Interestingly, the PRD region of tau can form clusters in cells when fused to a photosensitive oligomerization domain, unlike the MTBR region. Phospho-mimetic mutations (T231E, S235E or T231E/S235E) adjacent to the B1-1 epitope abolished light-induced phase separation.^25^ This suggests that phosphorylation in this region alters the charge distribution and affects the weak electrostatic interactions that mediate condensation. VHHs targeting the tau MTBR region do not robustly inhibit LLPS. This effect is observed more mildly with VHH Z70, but not at all for VHH A31, despite their binding to a similarly charged PHF6 (VQIVYK) and PHF6* (VQIINK) motifs, respectively. These results suggest that shielding charges alone are insufficient to explain the inhibitory effect. NMR binding studies indeed revealed that VHH B1-1 binding affects multiple resonances outside its epitope in full-length tau, but not in a fragment corresponding to the central region (PRD2, R1-R2, and part of R3). Although our data also indicate that electrostatic interactions play a dominant role in tau LLPS, the observation that VHH B1-1 binding affects resonances outside its epitope (as shown by NMR) suggests that conformational effects may also contribute to the modulation of tau condensation.

RNA acts as a scaffolding platform that brings tau molecules into close proximity. In the presence of VHH B1-1, LLPS of tau was observed at low RNA concentrations, but was abolished at higher concentrations. This may be because, at low RNA concentrations, the tau molecules remain close enough to undergo LLPS. However, at higher concentrations, the increased number of attachment points dilutes the tau-tau interactions, which prevents phase separation in the presence of VHH B1-1. VHH Z70 binds to the PHF6 sequence involved in tau nucleation and inhibits tau aggregation in the presence of heparin, as well as in cell and mouse seeding models.^19^ In this study, however, VHH Z70 lightly reduced tau LLPS in the presence of RNA, exhibiting a much weaker effect than VHH B1-1. Furthermore, VHH Z70 promoted tau LLPS in the presence of PEG. VHH B1-1, on the other hand, showed very low *in vitro* capacity to inhibit tau aggregation (Supplementary Figure 12). These results reinforce the idea that tau LLPS and cofactor-induced tau aggregation are distinct processes driven by different molecular determinants. Dissolution of tau droplets at high salt concentrations (300 mM NaCl) confirms that electrostatic interactions dominate tau LLPS under PEG- and RNA-inducing conditions. Our approach using VHH binding provides domain-resolved, direct evidence of the molecular determinants of tau LLPS, complementing deletion or mutation studies. We identified the PRD as a functional hotspot that mediates multivalent interactions during RNA-induced LLPS. Additionally, the specific effect observed for RNA-induced droplet formation, compared to PEG induced LLPS, suggests that the tau interaction network differs between the two types of LLPS, even though both are remaining predominantly driven by electrostatic interactions.

Whether regulating tau LLPS is a viable therapeutic strategy to counteract tauopathy development remains an open question at this stage. Tau LLPS and tau oligomerization or aggregation have been linked, and an increased potential for nucleation and seeding in the condensed tau state.^4,12,16,24^ This suggests the need to decrease tau LLPS in a pathological environment. On the other hand, studies have suggested that LLPS prevents the formation of oligomeric toxic tau species. TDP-43 and tau interaction promotes their co-condensation but also suppress tau fibril formation and seeding.^10,13^ This indicates conversely that stimulating the LLPS could have therapeutic value.

While our *in vitro* experiments provide novel insights, the cellular environment, with its complex array of binding partners, post-translational modifications, and crowding agents, may affect tau LLPS in ways that were not captured here. Importantly, VHHs offer molecular probes that can selectively modulate tau condensates in cells without disturbing MTBR-dependent functions, such as microtubule binding or aggregation. This novel approach may help answer the question of the link between tau LLPS and tauopathy development.

## Material and Methods

### Production and purification of tau2N4R and tauF3 and tauMTBD fragment

pET15b-tau2N4R, tau1N4RV5, tauF3 fragment (residues 208-324 of tau2N4R) or tauMTBD fragment (residues 245-369 of tau2N4R) recombinant T7 lac expression plasmids were transformed into competent *E. coli* BL21 (DE3) bacterial cells. A small-scale culture was then grown in LB medium at 37°C. To produce recombinant ^15^N-tau2N4R and ^15^N-tauF3, the small-scale culture was added to one liter of a modified M9 medium containing 1× MEM vitamin mix (Sigma-Aldrich), 4 g of glucose, 1 g of ^15^N-NH_4_Cl (Sigma-Aldrich), 0.5 g of ^15^N-enriched Isogro growth powder (Sigma-Aldrich), 0.1 mM CaCl_2_, and 2 mM MgSO_4_. To produce the other recombinant tau constructs, the small-scale culture was added to one liter of LB medium at 37°C.

Protein production was induced with 0.5 mM isopropyl-β-D-thiogalactopyranoside (IPTG) when the culture reached an optical density of 0.8 at 600 nm. The cells were lysed ina solution of 50 mM NaPi pH 6.5, 2.5 mM EDTA, containing one tab of complete protease inhibitor cocktail (Sigma-Aldrich). The tau proteins were first purified by heating the bacterial extract for 15 minutes at 75°C. After centrifugation, the resulting supernatant was then passed through a cation exchange chromatography resin (Hitrap SP Sepharose FF, 5 ml, Cytiva), which was equilibrated with 50 mM NaPi at pH 6.5. The tau proteins were eluted with a NaCl gradient. The tau proteins were the buffer-exchanged with 50 mM ammonium bicarbonate using a Hiload 16/60 desalting column (Cytiva) before lyophilization. Detailed procedures are available.^26,27^

### Selection and Screening of the VHHs Directed against tau protein

The Nali-H1 VHH library was screened against recombinant biotinylated tau.^28^ Recombinant tau protein was biotinylated using EZ-Link Sulfo-NHS-Biotin (Thermo Fisher Scientific), and the biotinylated tau protein bound to streptavidin-coated Dynabeads M-280 (Invitrogen) at concentrations that decreased gradually with each round of selection (100nM, 50nM and 10nM). A non-absorbed phage ELISA assay using avidin plates and biotinylated tau antigen (5 µg/ml) was performed as a second step of cross-validation. VHH Z70 was obtained from an additional optimization screen using the yeast two-hybrid method with tau protein as bait.^19,29^ VHH A31 was obtained from a separate phage display screening of the Nali-H1 library followed by a yeast two-hybrid-based optimization step.^19,29^

### Production and purification of VHHs

The VHHs were produced using a pET22b plasmid containing a recombinant sequence that encodes a pelB leader sequence, a 6-His tag, a tobacco etch virus (TEV) protease cleavage site, and the desired VHH, with or without an additional C-terminal cysteine.^29^ The VHH anti-V5 was developed to recognize the V5 tag, which is widely used and derived from a small epitope found on the P and V proteins of paramyxovirus in the simian virus 5 (SV5) family.^21^ Recombinant *E. coli* cells produced periplasmic proteins after induction with 1 mM IPTG in terrific broth medium. Production continued for four hours at 28°C or overnight at 16°C, after which cells were pelleted by centrifugation. The pellet was then resuspended in a solution of 200 mM Tris-HCl, 500 mM sucrose, 0.5 mM EDTA at pH 8. The solution was incubated on ice for 30 minutes. Then, the resuspended pellet was then diluted four times with water to yield a final concentration of 50 mM Tris-HCl, 125 mM sucrose, 0.125 mM EDTA at pH 8, and complete protease inhibitor (Roche). Incubation continued on ice for another 30 minutes. After centrifugation, the supernatant corresponding to the periplasmic extract was recovered. The VHHs were purified by immobilized-metal affinity chromatography (IMAC) (HisTrap HP, 1 mL, Cytiva), followed by size exclusion chromatography (HiLoad 16/60 Superdex 75, prep grade, Cytiva) in a phosphate buffer solution containing 50 mM sodium phosphate buffer [NaPi], pH 6.7, 30 mM NaCl, 2.5 mM EDTA, and 1 mM dithiothreitol (DTT). The VHHs were then dialyzed against 50 mM Tris at pH 8, 50 mM NaCl, and cleaved with His-tagged Tobacco Etch Virus (TEV) protease. A second IMAC step removed the TEV protease and the cleaved 6-His tag, and the VHHs were recovered in the flow-through. The VHHs were then concentrated and flash-frozen for further use.

### Characterization of the interactions between tau, tau fragments and the VHHs

Affinity measurements were performed using a Biacore T200 optical biosensor instrument (Cytiva). The recombinant tau2N4R and tau1N4RV5 proteins were biotinylated using a 3-fold molar excess of NHS-biotin conjugates (Thermo Fisher Scientific) for 4 hours at 4 °C. The recombinant tauF3 and tauMTBD proteins were biotinylated using a 1 to 1 molar ratio with NHS-biotin conjugates (Thermo Fisher Scientific) for 4 hours at 4 °C. The reactions were then quenched with 10 mM Tris-HCl at pH 7.4. Residual NHS-biotin was removed by performing two consecutive buffer exchanges using a Zeba Spin Desalting Column (Thermo Fisher Scientific). Capture of biotinylated tau proteins in HBS-EP+ buffer (Cytiva) was performed on SA sensor chips (Cytiva), until the total amount of captured proteins reached 300 resonance units for each. A free flow cell was always used as a reference to assess nonspecific binding and to allow for background correction. VHH anti-V5 was injected sequentially at increasing concentrations ranging from 0.0625 to 4 μM in a single-cycle at a flow rate of 30 µL/min. VHH B1-1 was injected sequentially at increasing concentrations ranging from 0.125 to 2 μM in a single-cycle at a flow rate of 30 µL/min. Three consecutive 1 M NaCl washes were performed for regeneration between each VHH injection. Single-cycle kinetic analysis was performed to determine the association k_on_ and dissociation k_off_ rate constants, as well as the dissociation equilibrium constant K_D_, by curve fitting of the sensorgrams using the 1:1 Langmuir interaction model in BIAevaluation software 2.0 (Cytiva).^30^

### Characterization of the interaction of VHH B1-1 with tau epitope-derived peptide

Affinity measurements were performed using a BIAcore T200 optical biosensor instrument (Cytiva). Recombinant VHH B1-1, with a cysteine at its C-terminus, was biotinylated with 10-fold molar excess of maleimide biotin conjugates (Thermo Fisher Scientific) overnight at 4°C. The reaction was then quenched with 2 mM DTT. Residual NHS-biotin was removed by performing two consecutive buffer exchanges using a Zeba spin desalting column (Thermo Fisher Scientific).

Biotinylated VHH capture was performed on a SA sensor chip in HBS-EP+, as was done for tau immobilization. A flow cell was used as a reference to assess nonspecific binding for background correction. Biotinylated VHH B1-1 was injected onto an SA chip until the total amount of captured proteins reached 660 RU. Nine consecutive and increasing ½ log concentrations of the tau peptide _221_REPKKVAVVRTP_232_ were injected sequentially at a flow rate of 30 µL/min, in a multi-cycle, with regeneration (one wash with 1 M NaCl between each peptide injection). K_D_ determination was obtained by plotting the individual response value for each tau peptide concentration. Equilibrium data were fitted to a binding model using non-linear regression analysis (one binding site, specific) available in GraphPad Prism software. The stoichiometric ratio (SR) value is the ratio of the observed maximum binding response to the theoretical maximum binding response (Theo Rmax). Theoretical Rmax is calculated using the molecular weight (MW) ratio of the analyte to the ligand, multiplied by the expected stoichiometry and the amount of immobilized ligand (in RU), as follow:

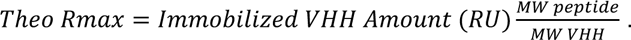

### NMR spectroscopy experiments

The analysis of the ^15^N-tau and ^15^N-tauF3/VHH B1-1 interactions was performed at 293 K using a Bruker Avance Neo 900 MHz spectrometer equipped with a cryogenic probe. Trimethylsilyl propionate was used as the internal proton reference. Lyophilized ^15^N tau was dissolved in an NMR buffer solution containing 10% D_2_O and mixed with a VHH at a final concentration of 100 μM for each protein. Two hundred microliters of each mixture in 3-mm tubes was sufficient to obtain the 2D ^1^H, ^15^N heteronuclear single quantum coherence (HSQC) spectra with 32 scans. The ^1^H, ^15^N HSQC spectra were acquired with 3072 and 416 points in the direct and indirect dimensions for spectral windows of 12.6 and 25 ppm, in the ^1^H and ^15^N dimensions, respectively. The data were processed with Bruker Topspin 3.6 (https://www.bruker.com/en/products-and-solutions/mr/nmr-software/topspin.html) and analyzed with POKY (poky.clas.ucdenver.edu).^31^ The spectra were rendered using POKY, and the intensity plots were designed using matplotlib.^32^

### Tau fragment library screening

The construction of the tau Fragment library is described.^18^ The VHH B1-1 coding sequence was amplified by PCR and cloned into pB29 as an N-terminal fusion to LexA (VHH B1-1_LexA). This construct was then used to produce a bait for screening the tau fragment library, which was constructed into pP7. The tau fragment library was screened using a mating approach with the yeast strains YHGX13 (Y187 ade2-101::loxP-kanMX-loxP, matα) and L40ΔGal4 (mata).^33^ Twenty-four His^+^ colonies, corresponding to 535000 tested diploids, were selected on a medium lacking tryptophan, leucine, and histidine and supplemented with 0.5 mM of 3-Aminotriazole. The cDNA of the prey fragments from the positive clones was amplified by PCR and sequenced at the 5’ and 3’ junctions.

### Turbidity assays

When PEG (10kDa, Sigma) was used as an inducer, a Tris 50mM, NaCl 10mM at pH 7,4 buffer solution was used. Tau and VHHs were added to the samples at equimolar concentrations of 10 µM, and LLPS formation was induced by adding PEG at a final concentration of 7% in the presence of 1 mM DTT. The final concentration of NaCl was 300mM when added. For phase diagram experiments, the tau protein concentration increased from 0 µM to 10 µM in increments of 2 μM, and the PEG concentration increased from 0 to 14 % in increments of 2 %. The F8-2 VHH concentration was set to 10 µM and did not vary.

When using RNA polyA as an inducer (Sigma), HEPES (25 mM), NaCl (10 mM) pH 7.4 were used. The solubilized 2N4R Tau was then centrifuged at 16,000 g for 10 minutes to remove any gel that may have formed during resuspension. Tau and VHHs were added to the samples at equimolar concentrations of 20 µM, and LLPS formation was induced by adding RNA at a final concentration of 50 µg/mL in the presence of 1 mM DTT. The final concentration of NaCl was 300 mM when added. Optical density tests were carried out in 96-well plates with a sample volume of 90 µL. The optical density at 400 nm was measured using a Pherastar FS plate reader (BMG Labtech) at room temperature . OD_400_ values were normalized to the tau LLPS control conditions to take into account differences of absolute OD_400_ values that appear between the different tau production batches .

For phase diagram experiments, the tau protein concentration increased from 0 µM to 20 µM in increments of 4 µM, and the RNA concentration increased from 0 µg/mL to 125 µg/mL in increments of 25 µg/mL. The B1-1 VHH concentration was set to 20 µM and did not vary.

### Dynamic Light Scattering

Sample preparation for dynamic light scattering was similar to that for the optical density assay. The same buffers were used in both processes, and equimolar concentrations of tau and VHH were employed. All reagents were centrifuged for 10 minutes at 21,000 g and 4 °C to pellet and remove potential impurities.

When PEG was used as the inducer, the tau and VHH concentrations were 10 µM, and polyethylene glycol was added at a final concentration of 7% in the presence of 1 mM DTT. When using RNA as the inducer, the tau and VHH concentrations were 20 µM, and the RNA was added at a final concentration of 50 µg/mL in the presence of 1 mM DTT. The final NaCl concentration was 300 mM when added. The samples were produced fresh and the inducer was added directly to the quartz cuvette used for the measurement. Then, the sample was mixed by pipetting up and down. Measurements were performed on a Zetasizer Nano-S (Malvern Instruments) at 25°C with a scattering angle of 173°. Ten measurements were performed for each sample, and it was ensured they comply with the autocorrelation. All conditions were repeated at least three times. For the analysis, the viscosity was adjusted to 0.8703 cP for all samples, and the hydrodynamic radii were calculated using the Stokes–Einstein equation.

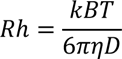

### Fluorescent labeling of recombinant tau2N4R and VHHs

Solubilized tau in HEPES was centrifuged at 16 000g for 10minutes to remove the potential gel formed by the resuspension. The recombinant tau2N4R was labelled using a 3-fold molar excess of Maleimid-TAMRA (Euromedex) for 2 hours at room temperature. to attach the dye to its two native cysteine residues. The recombinant VHH-cystein were labelled using a 1 to 1 molar ratio with Maleimid Atto488 (Sigma) for 2 hours at 4°C.The reactions were then quenched with 2 mM DTT. Residual probes were removed by two consecutive buffer exchanges using a Zeba spin desalting column (Thermo Fisher Scientific).

### Confocal Microscopy analysis

Microscopy slides (18-well µslides, Ibidi) were coated with a coating solution overnight at 4°C. Coating solution is composed of Laminin (10µg/mL, Sigma) and poly-D Lysin (0.5mg/mL, Sigma). Tau LLPS were generated using the same procedures as turbidity and DLS assays. Samples were freshly produced before depositing 15µL on the slide. The acquisition of the images was done with the confocal microscope (ZEISS 980 Airyscan). Three independent experiment corresponding for each to nine different images were recorded using a 63x oil objective. For the samples produced with PEG, three optical fields were used. For the sample produced with RNA, the optical field used was close to the bottom of the slides as the LLPS sedimented faster. The analysis of the images was automatized with a script on the FIJI: ImageJ software to quantify the labelled surface area of condensates on each image.

### Fluorescence Recovery After Photobleaching

Tau-VHH LLPS containing labeled tau-TAMRA were imaged before and directly after bleaching with a 561 nm laser. At ×63 magnification and ×4 optical zooming, droplets were outlined with a circular “bleaching” overlay object. The FRAP parameters were as follows: 50 pre-bleaching images, bleaching (laser power 100% intensity; 10 iterations), an immediate post-bleach image and several images over a total of 200 s (Figures display T15, T16 and T200 second images). The droplet region was manually segmented and the average fluorescence of the bleached region in the droplet I_rel_ at time t after bleaching was measured to correct for photobleaching during acquisition, as described previously.^34^ The FRAP curves were then normalized by setting the pre-bleach fluorescence intensity values to one and the intensity immediately after bleach to zero. All images were acquired with the same settings (i.e. scan speed, resolution, magnification, optical zoom, gain, offset and laser intensity). The FRAP experiments were repeated two independent times with the intensities from all droplets analyzed in each experiment averaged together.

### *In vitro* kinetic tau aggregation assay

Tau 2N4R aggregation assays were performed with 10 μM Tau and with increasing concentrations of VHH B1-1 (between 0 and 10 μM) in buffer containing 50 mM MES pH 6.9, 30 mM NaCl, 2.5 mM EDTA, 0.3 mM freshly prepared DTT, 2.5 mM heparin H3 (Sigma-Aldrich), and 50 mM Thioflavin T (Sigma-Aldrich), at 37°C. The resulting fluorescence of Thioflavin T was recorded every 5 min/cycle within 75 cycles using PHERAstar microplate-reader (BMG labtech). The measures were normalized in fluorescence percentage, 100% being defined as the maximum value reached in the positive Tau control.

## Acknowledgments

We thank the BiCel platform of US 41 - UAR 2014 - PLBS for the microscopy data acquisition. The NMR facilities were funded by the Conseil Régional du Nord, the CNRS, the Institut Pasteur de Lille, the European Union, the French Ministry of Research and the University of Lille. Financial support from the IR INFRANALYTICS FR2054 CNRS for the conduct of the research is gratefully acknowledged. We are grateful to Dr JeanChristophe Rain (Hybrigenics Services, FR) for providing the plasmid anti-V5 VHH PHEN2. We particularly thank Dr Guilherme. G. Moreira and Professor Cláudio M. Gomes (Universidade de Lisboa) for their valuable advices and inputs for tau LLPS characterization at the start of this project.

## Funding

This project was funded by the France Alzheimer’s Association (Project NanoLIPS 6433 to C.D.) This project benefited from financial contribution from the Agence Nationale de la Recherche, France (projects ANR-11LABX-01 to L.B. and I.L., including S.T. fellowship; ANR-18-CE44-0016 to L.B. and I.L.; and ANR-22CE92-0061 to L.B. and I.L., including C.D. fellowship), the European Union (HORIZON-MSCA-2022-DN-01, Project TAME, GA101119596, including E.M. fellowship) and the Rainwater Charitable Foundation & Alzheimer’s Association (Project T-PEP-23-969176, including C.D. fellowship). We also thank the *Fondation pour la Recherche Médicale* (project SMC202405019700 to D.B.).

## Author contributions

Investigation: S.T, L.M, E.M, M.N, J.M, L.H; F.-X.C, C.D. Methodology: S.T, E.D, C.D. Visualization: E.D, C.D. Formal analysis: S.T, E.D, C.D. Funding Acquisition: D.B., L.B, I.L, C.D. Supervision: V.B-S, I.L, C.D. Conceptualization: C.D. Project administration: C.D. Writing – Original Draft: S.T., I.L., C.D. Writing – Review & Editing: All authors.

## Competing interests

C.D., E.D., L.B., and I.L. are the inventors of a patent (WO2020/120644A1) that covers the use of VHH Z70. The remaining authors declare no competing interests.

