## Supplementary Information for "A proline-rich-domain-binding single domain antibody selectively inhibits RNA-induced liquid-liquid phase separation of tau"

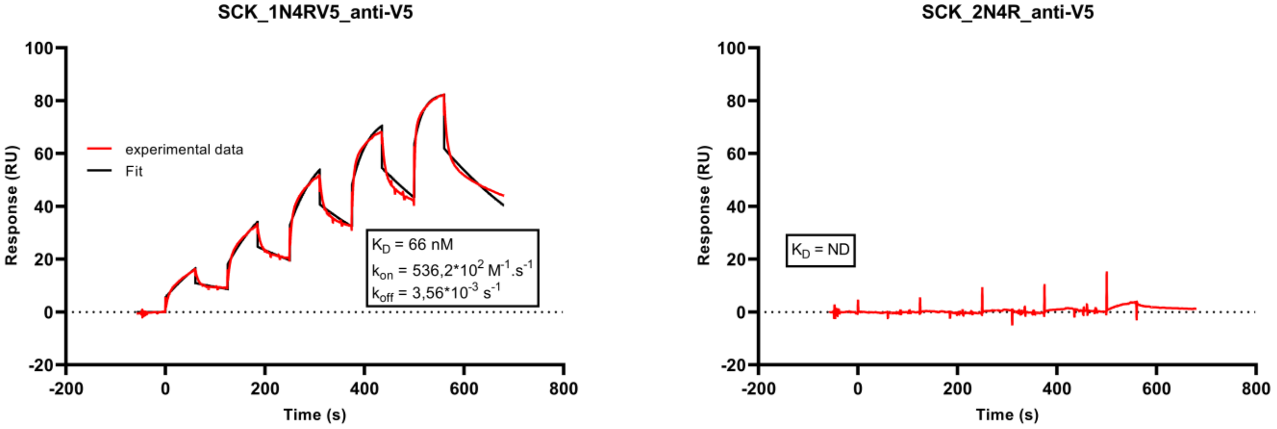


**Supplementary Figure 1: VHH anti-V5 binds specifically to tau1N4RV5 with a $K_D$ in the sub-micromolar range.**

We tested VHH anti-V5 as an analyte using single-cycle kinetics (SCK). SCK consists of increasing the analyte concentration in five consecutive injections with a short dissociation time and no regeneration between injections. We measured the affinity of the interaction between a recombinant tau version in which we inserted exon 2 of the longest tau isoform's coding sequence into the V5 tag (1N4RV5) and VHH anti-V5. We obtained a KD of 66nM, which is consistent with the previously published KD of 29nM determined by a FACS experiment. Using the same procedure, we confirmed that VHH anti-V5 does not bind to tau2N4R.

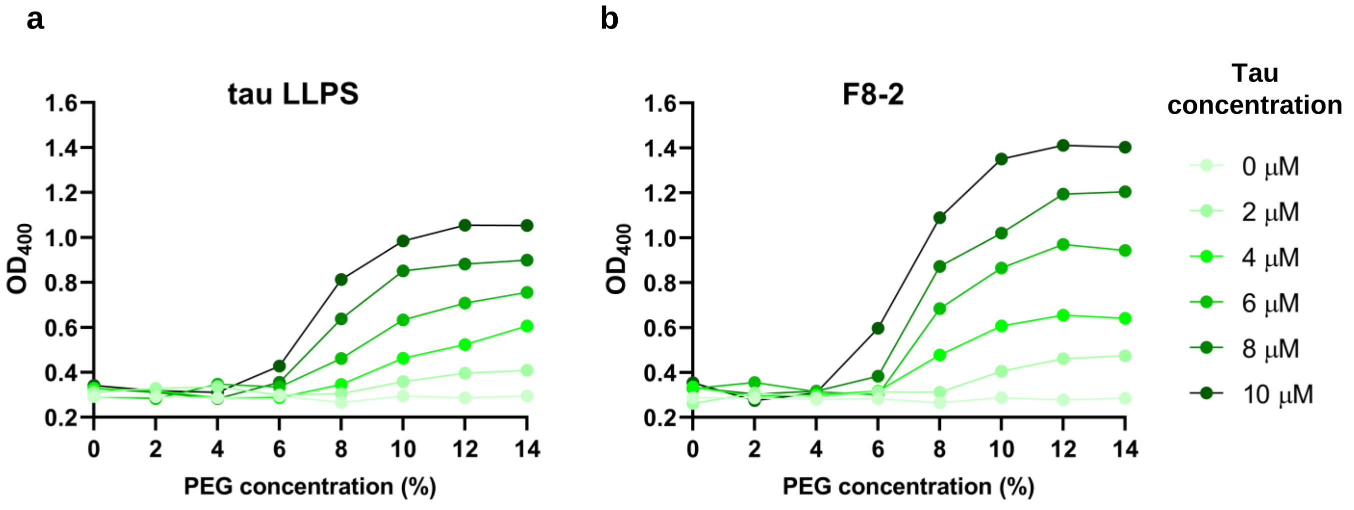


**Supplementary Figure 2a-b: Phase diagram results for tau2N4R phase separation in the presence of PEG and VHH F8-2.** The plot shows PEG-induced LLPS followed by turbidity (optical density at 400 nm) at increasing tau2N4R concentrations ranging from 0 to 10 μM and increasing PEG concentrations up to 14%. This is shown in the absence (a) or presence (b) of a fixed concentration of VHH F8-2 (10 μM). Averaged values corresponding to two replicates of tau2N4R droplet preparation are shown for each condition tested.

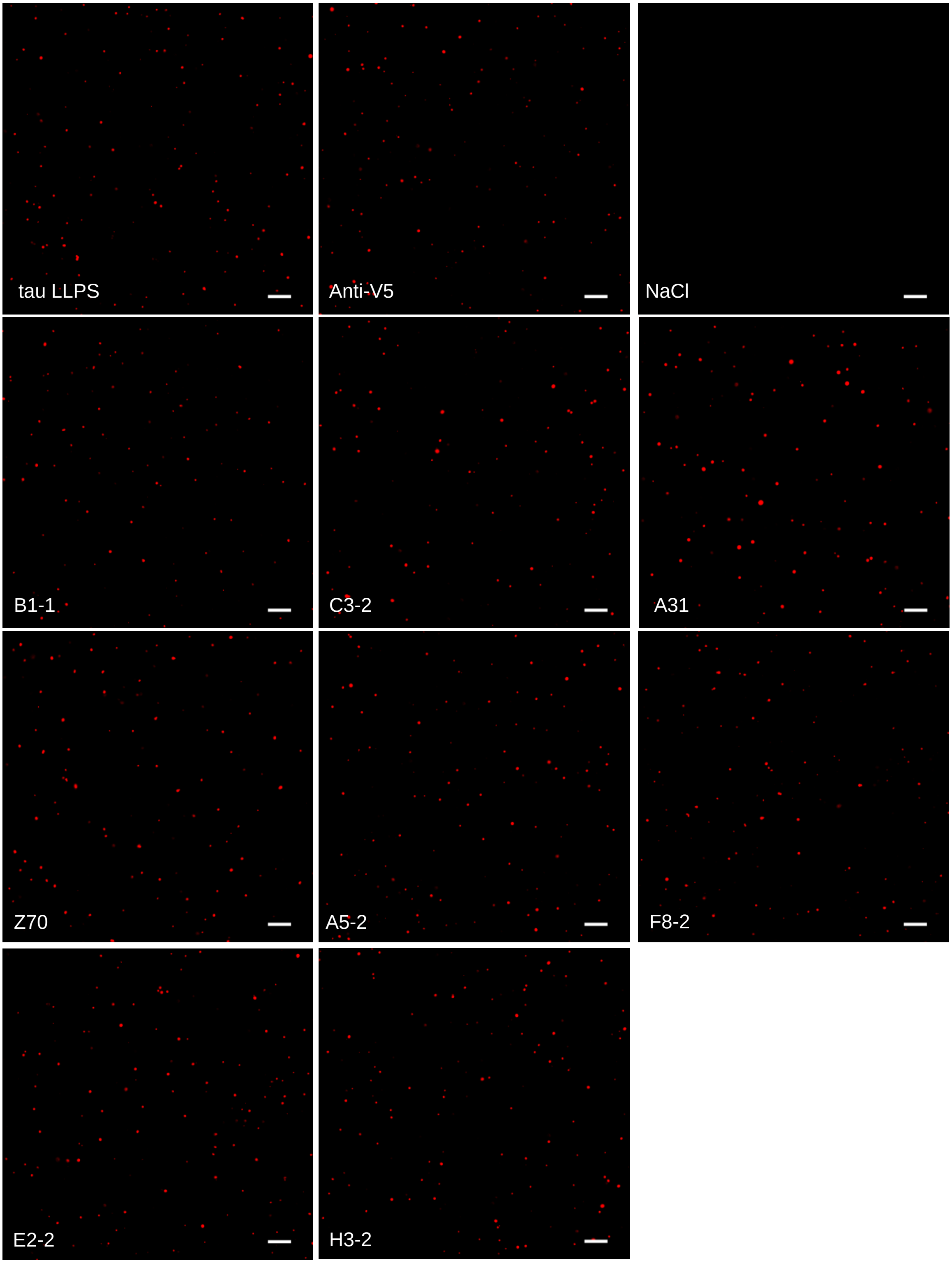


**Supplementary Figure 3: Visualization of tau LLPS formation using confocal microscopy in the presence of anti-tau VHHs and 7% PEG.** Confocal microscopy images of tau LLPS formation were obtained using 10 µM tau-TAMRA, which was co-incubated with 10 µM VHH (anti-V5 or anti-tau) and 7% PEG. The images are representative of three independent experiments per condition. Tau was visualized in red. The scale bar is 10 µm.

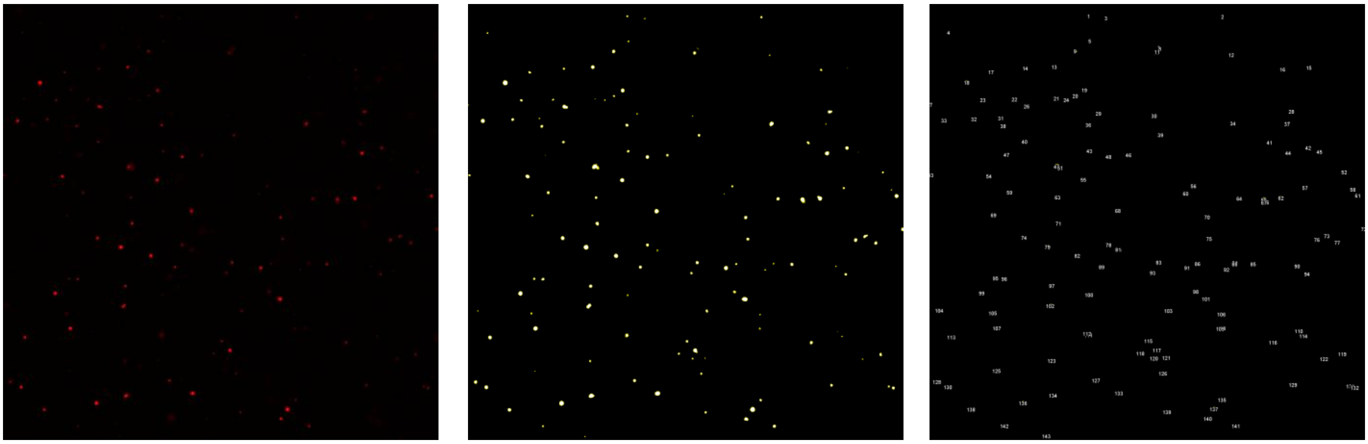

**Supplementary Figure 4: Quantification strategy for characterizing tau droplets**
Modulation of tau LLPS formation by the VHHs was characterized using the following described strategy. Tau LLPS formation was visualized with a 63x objective and a confocal microscope (left image). Nine images were recorded per experiment (three independent experiments were performed per condition). A fluorescence filter was set to remove the fluorescence background from the images and isolate the fluorescence signal from the tau LLPS (middle image). Then, the fluorescence surface area of each individual event was quantified (right image), and the total surface area was calculated.

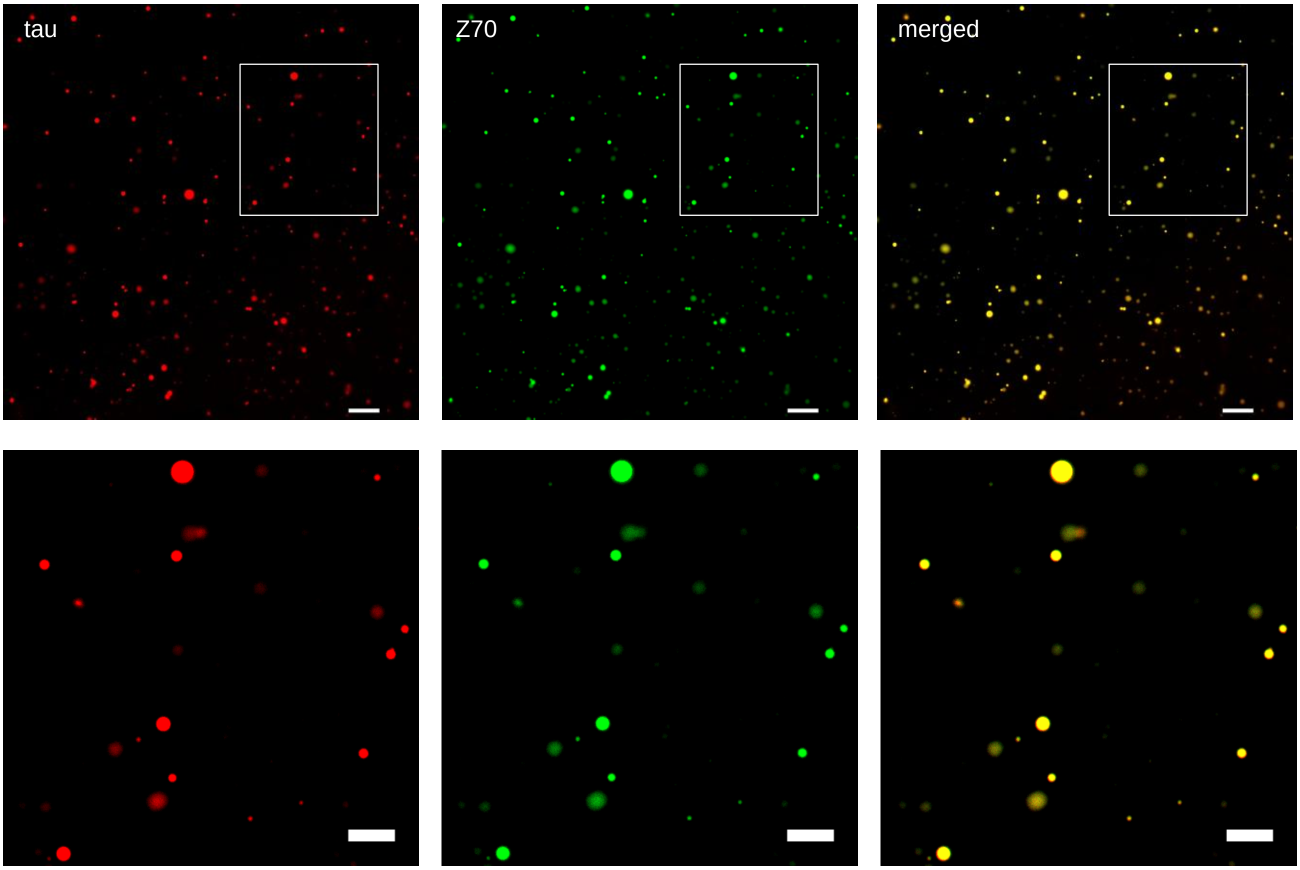


**Supplementary Figure 5 : The anti-tau VHH Z70 protein is located inside tau droplets.** Representative fluorescence images, and associated zoom images, of tau LLPS formed in the presence of 10 µM tau-TAMRA, which was co-incubated with 10 µM VHH Z70-Atto488 and 7% PEG. The scale bars are 10 µm and 5 µm, respectively.

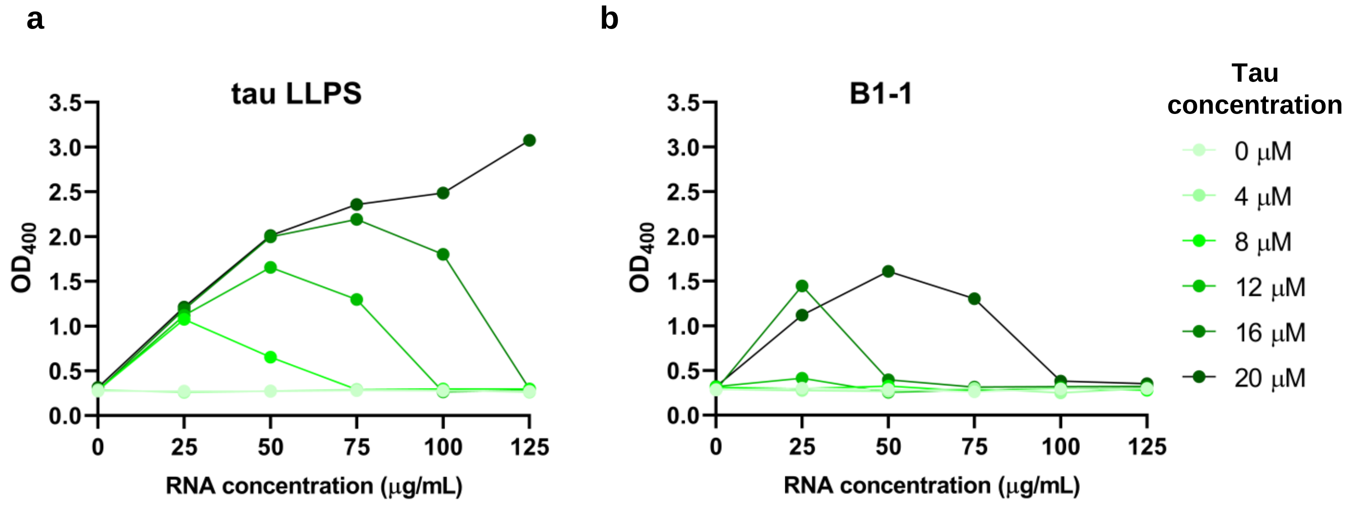


**Supplementary Figure 6a-b: Phase diagram results for tau2N4R LLPS formation in the presence of RNA polyA and VHH B1-1.** The plot show RNA-induced LLPS followed by turbidity (optical density at 400 nm) at increasing tau2N4R concentrations from 0 to 20 μM and increasing RNA polyA concentrations up to 125 μg/mL, in the absence (a) or presence (b) of a fixed VHH B1-1 concentration of 20 μM. The averaged values correspond to two replicates of tau2N4R droplet preparation for each condition tested.

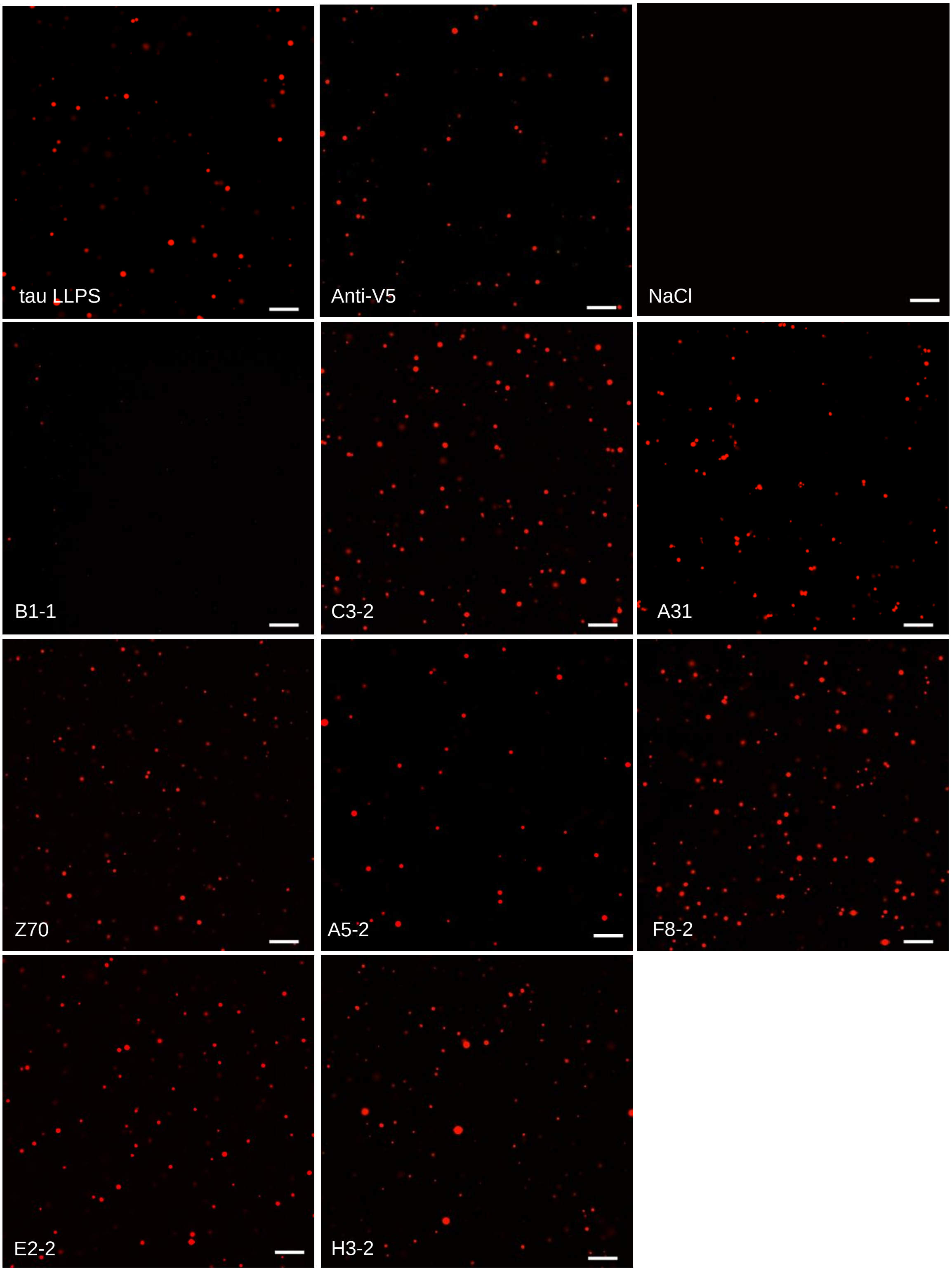


**Supplementary Figure 7: Confocal microscopy visualization of tau LLPS formation in the presence of anti-tau VHHs and 50 µg/mL of RNA polyA.** Confocal microscopy images of tau LLPS formation were obtained using 20 µM tau-TAMRA, which was co-incubated with 20 µM VHH (anti-V5 or anti-tau). Images are representative of three independent experiments per condition. Tau was visualized in red. The scale bar is 20 µm.

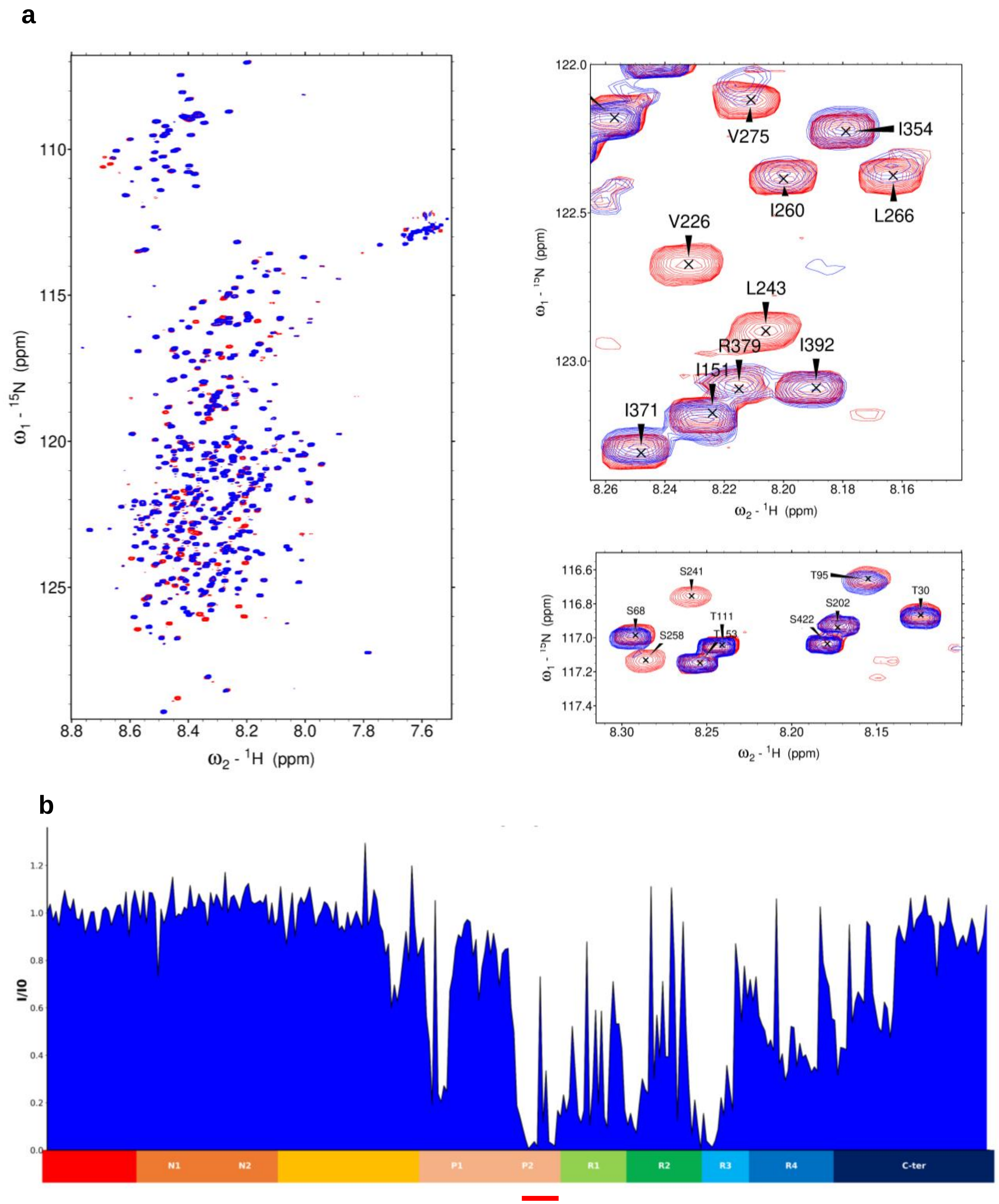


**Supplementary Figure 8: VHH B1-1 binds to the proline-rich domain of tau**

**a**. Overlay of two-dimensional $^{1}H$, $^{15}N$ HSQC full spectra and details of free full-length tau2N4R (in red) and tau2N4R mixed with unlabeled VHH B1-1 (in blue). The spectra are representative of two independent experiments. In the spectrum of tau with VHH B1-1, several resonances corresponding to the different tau regions are broadened beyond detection compared to the tau control spectrum. **b**. Normalized NMR intensities (I/I0) along the tau2N4R sequence. I0 and I correspond to the resonance intensity when tau is free in solution or mixed with an equimolar amount of VHH B1-1 (I), respectively. Plotting the normalized intensity ratio (I/I0) allowed the identification of the proline-rich domain sequence as the target of VHH B1-1 interaction. A red line indicates the region containing the corresponding major broadened resonances. N1 is a sequence in the N-terminal domain that is not present in all tau isoforms (named tau 0N, tau 1N or tau 2N). The proline-rich domain is subdivided in P1 and P2 regions. The MTBD consists of four partially repeated regions, R1 to R4.

**a**

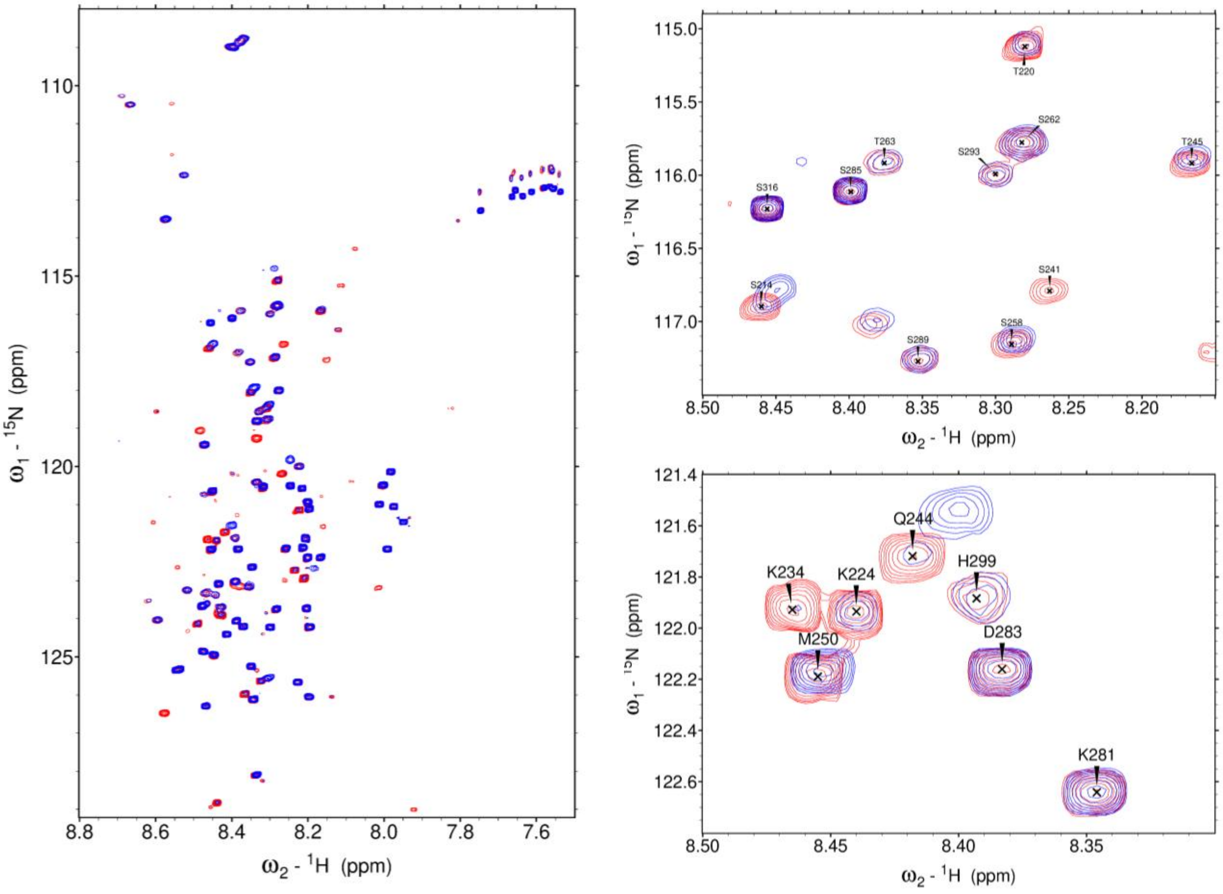


**b**

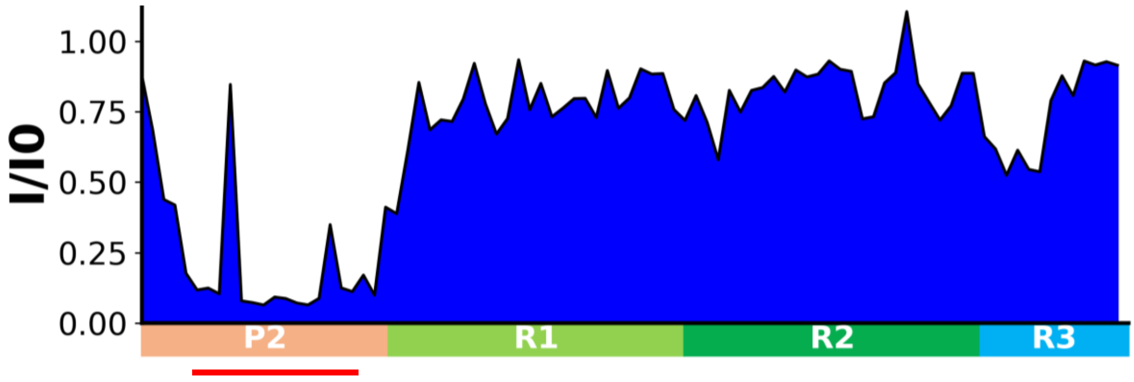


**Supplementary Figure 9: 2D HSQC NMR experiment of tauF3 fragment identified the tau sequence 220-243 as central to tau-B1-1 binding**

**a**. Overlay of two-dimensional $^{1}H$, $^{15}N$ HSQC full spectra and details of free tauF3 fragment, residues 208-334, in red and the tauF3 fragment mixed with unlabeled VHH B1-1, in blue. The spectra are representative of two independent experiments. The spectrum of tauF3 in the presence of VHH B1-1 showed that resonances corresponding to the proline-rich domain were broadened beyond detection compared to the tauF3 control spectrum. **b**. Normalized NMR intensities (I/I0) along the tauF3 sequence. I0 and I correspond to the resonance intensity when tau is free in solution or mixed with an equimolar amount of VHH B1-1 (I), respectively. The normalized intensity ratio plot (I/I0) allowed the identification of the proline-rich domain sequence as the target of the VHH B1-1 interaction. A red line indicates the region containing the corresponding major broadened resonances.

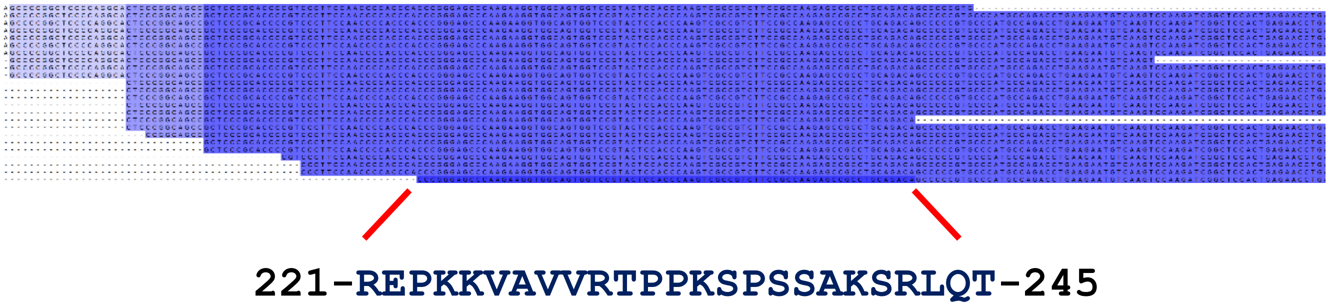


**Supplementary Figure 10: Identification of the minimal epitope recognized by VHH B1-1 using tau fragment library and yeast two hybrid.**

Sequence alignment of the 24 Tau fragments corresponding to the 24 positive colonies picked on selective growth conditions and thus binding VHH B1-1. The minimal common sequence is highlighted. Sequences are not meant to be read.

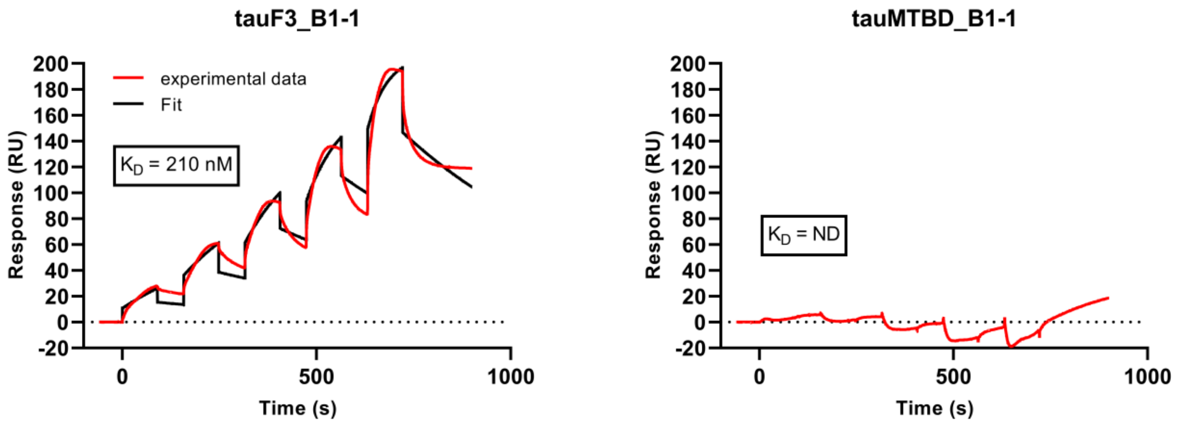


**Supplementary Figure 11: VHH B1-1 binds specifically to the proline-rich domain of tau**
Sensorgrams (reference-subtracted data) of single cycle kinetics (SCK) analysis performed on immobilized biotinylated tauF3 and biotinylated tauMTBD fragments (residues 245-369), with five injections of increasing concentrations of VHH B1-1. The sensorgrams are representative of two independent experiments. $K_D$ values are included in the sensorgram. The black lines correspond to the fitted curves and the red lines correspond to the measurements.

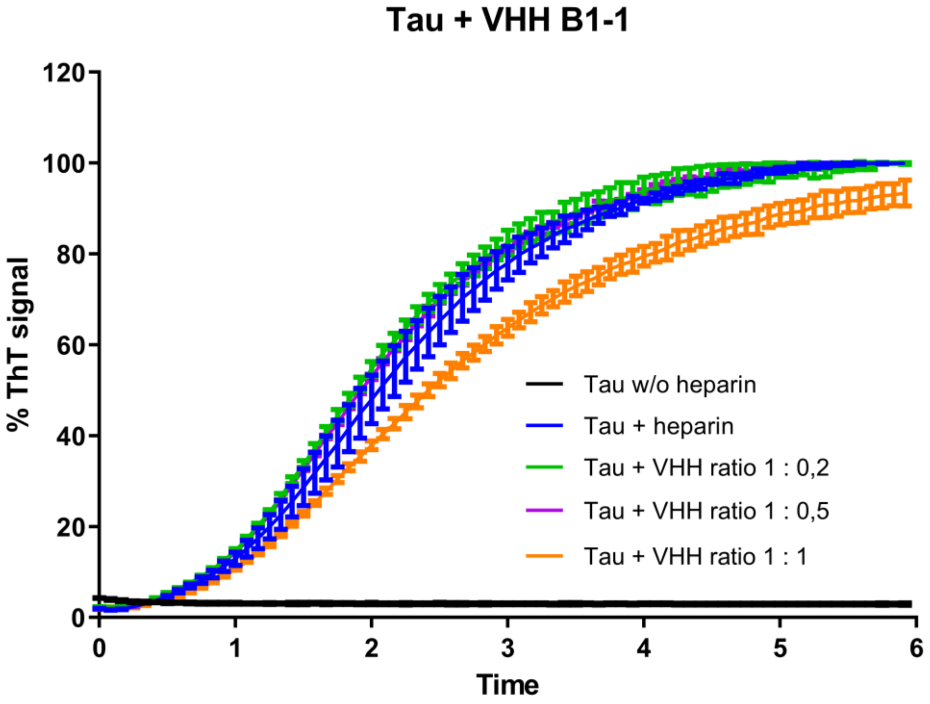


**Supplementary Figure 12: VHH B1-1 does not strongly inhibit tau aggregation in vitro**
Aggregation of Tau (10 µM) in the absence of heparin (black curve), in the presence of heparin (blue curve) and of increasing concentration VHH B1-1 (2, 5, and 10 µM) monitored by Thioflavin T fluorescence at 490 nm (n = 1 performed in triplicate). Error bars: SD.
